# Respiratory complex III_2_ assembles complex I via toxic intermediate in mitochondrial disease

**DOI:** 10.1101/2025.06.17.660237

**Authors:** Maria G. Ayala-Hernandez, Anetzy Bermudez Torales, Hannah Camille Tan, Claire B. Montgomery, Abhilash Padavannil, Gino Cortopassi, James A. Letts

**Affiliations:** Department of Molecular and Cellular Biology, University of California, Davis, United States; Department of Molecular Biosciences, School of Veterinary Medicine, University of California, Davis, United States

## Abstract

Mutations in mitochondrial complex I can cause severe metabolic disease. Although no treatments are available for complex I deficiencies, chronic hypoxia improves lifespan and function in a mouse model of the severe mitochondrial disease Leigh syndrome caused by mutation of complex I subunit NDUFS4. To understand the molecular mechanism of NDUFS4 mutant pathophysiology and hypoxia rescue, we investigated the structure of complex I in respiratory supercomplexes isolated from NDUFS4 mutant mice. We identified complex I assembly intermediates bound to complex III_2_, proving the cooperative assembly model. Further, an accumulated complex I intermediate is structurally consistent with pathological oxygen-dependent reverse electron transfer, revealing unanticipated pathophysiology and hypoxia rescue mechanisms. Thus, the build-up of toxic intermediates and not simply decreases in complex I levels underlie mitochondrial disease.

## Main Text

Mitochondria provide the energy needed to drive mammalian biochemistry, powering muscle contraction, neuronal signaling, and anabolism (*1–4*). The respiratory electron transport chain (ETC) in the inner mitochondrial membrane (IMM) is responsible for converting energy ingested as food into a proton electrochemical gradient that can be used for the synthesis of adenosine triphosphate (ATP) by the ATP synthase complex. The ETC is composed of three large proton-pumping membrane protein complexes: complex I (CI), a proton-pumping NADH:ubiquinone (CoQ) oxidoreductase; complex III (CIII), a proton-pumping ubiquinol (CoQH_2_):cytochrome c (cyt *c*) oxidoreductase; and complex IV (CIV) a proton-pumping cytochrome *c* oxidase. CI, an obligately dimeric CIII (CIII_2_) and/or CIV can assemble within the IMM to form higher order structures known as supercomplexes (SCs). Although, SCs can vary in composition and abundance across organisms and tissues, SC I+III_2_ (SC_1,3_) and SC I+III_2_+IV (known as the respirasome, R), are prevalent in mammalian mitochondria (*5, 6*). Cryogenic electron tomography (cryoET) and single particle analysis of isolated bovine heart mitochondria have shown that most CI is present in SCs in a structurally conserved association with CIII_2_ (*5, 7*). Mammalian CI is composed of 45 protein subunits organized into an electron-transferring peripheral arm (PA) that extends into the mitochondrial matrix and a proton-pumping membrane arm (MA) imbedded in the IMM (Fig. S1A). The peripheral arm is formed by two functional modules: the N-module, which accepts electrons from the reduced form of nicotinamide adenine dinucleotide (NADH), and the Q-module, which is adjacent to membrane and receives the electrons for CoQ reduction (Fig. S1A) (*8*). Assembly of CI proceeds via modules (Fig. S1B) and it is debated whether complexes must be fully assembled before associating (assembly first plasticity model) (*9*), or if assembly intermediates associate with CIII_2_ and complete assembly as part of a SC (cooperative assembly model; Fig. S1B) (*10–13*). CI and SC assembly defects have been proposed to underlie mitochondrial disease pathophysiology (*14, 15*), making them essential processes to fully understand.

Approximately one third of ETC diseases result from deficiencies in CI (*16–18*). In mammals, CI is composed of 14 highly conserved core catalytic subunits and 31 accessory subunits (*19*). Knockout (KO) studies and naturally occurring mutations show that disruption of the accessory subunit NDUFS4 (Fig. S1C) leads to the severe multisystemic progressive neurodegenerative disorder Leigh syndrome (*20*). Most Leigh syndrome patients die by the age of two and no treatments are currently available (*21, 22*). The current best-characterized model of mitochondrial disease in mammals is the Palmiter NDUFS4 knockout mouse (S4^KO^), which experiences loss of hair and vision, ataxia, and death before reaching adulthood (*20, 23*). S4^KO^ mice show significant CI deficiency but retain some CI activity (*24*). Studies from S4^KO^ mice and Leigh syndrome patients show that all intact CI is found in SCs and that partially assembled CI accumulates in SCs as well (*14, 25, 26*). This led to the hypothesis that in patients and mice with NDUFS4 disruption, CI assembly is rescued by SC formation (*14, 27*). Further, SC formation was recently shown to mask CIII_2_ deficiencies in another mouse model (*28*), suggesting a general role for SCs in mitigating ETC deficiency. In landmark papers, the Mootha and Zapol labs demonstrated that chronic hypoxia attenuates mitochondrial disease caused by the S4^KO^ in mice (*29, 30*). More recently, Meisel *et al.* demonstrated that hypoxic rescue of S4^KO^ is evolutionarily conserved in the nematode *Caenorhabditis elegans* and other mutations in CI subunits NDUFS7 (*nduf-7(et19)*) and NDUFS2 (*gas-1(fc21)*) can also be partially rescued by hypoxia (*31*). Further, they found that the secondary mutations G60D in the NDUFA6 subunit (NUDFA6^G60D^; NUO-3 in *C. elegans*) and R126Q in the NDUFA5 subunit (NDUFA5^R126Q^) phenocopy hypoxia rescue (*31*). Functional characterization of the mutants led Meisel *et al.* to conclude that although hypoxia led to increased CI content in both S4^KO^ mice and NDUFS2/*gas-1(fc21)* worms, increasing forward electron transport (FET) from NADH to CoQ was sufficient to rescue (*31*). However, the mechanism by which hypoxia and NDUFA6^G60D^ support CI FET, why enhancing CI FET alone is sufficient to rescue the respiratory deficiency and whether this mechanism is conserved across the three CI deficient mutants remains unknown. Thus, a major gap remains in our understanding of the pathophysiology of Leigh syndrome and other mitochondrial disorders of the ETC. Given the accumulation of CI assembly intermediates in the S4^KO^ mouse (*14*), understanding the pathway of CI assembly is needed to fully understand how its disruption may contribute to Leigh syndrome pathophysiology. Thus, we set out to biochemically and structurally characterize SCs containing CI intermediates from S4^KO^ mouse liver mitochondria.

### Fully assembled CI and CI assembly intermediates are present in SCs

Mitochondria were isolated from WT and S4^KO^ C57BL/6 murine livers; membrane protein complexes were extracted using the mild detergent digitonin and partially purified using a size exclusion chromatography (SEC) column (Fig. S2A, 2B). Fractions from the SEC containing SCs were pooled, concentrated and used for cryoEM grid preparation (Fig. S2C). After initial optimization, this resulted in sample-freezing less than nine hours after extraction. To confirm that CI from the S4^KO^ samples was not degrading on this timescale, we determined a time course of rotenone-inhibited NADH:CoQ activity in the detergent-extracted samples (Fig. 1A). The half-life of CI activity was 385 hours (291-551 hours 95% confidence interval) for the wild-type sample on ice in digitonin and 145 hours (120-181 hours 95% confidence interval) for the S4^KO^ sample in equivalent conditions (Fig. 1A). When the S4^KO^ mitochondrial membranes were extracted with the harsher detergent dodecyl-maltoside (DDM), the half-life of CI activity decreased to 37 hours (30-46 hours 95% confidence interval; Fig. 1A). Overall, these results demonstrated that CI was less stable in the S4^KO^, but sufficiently stable in our conditions that degradation products should not accumulate over nine hours.

**Figure 1.**
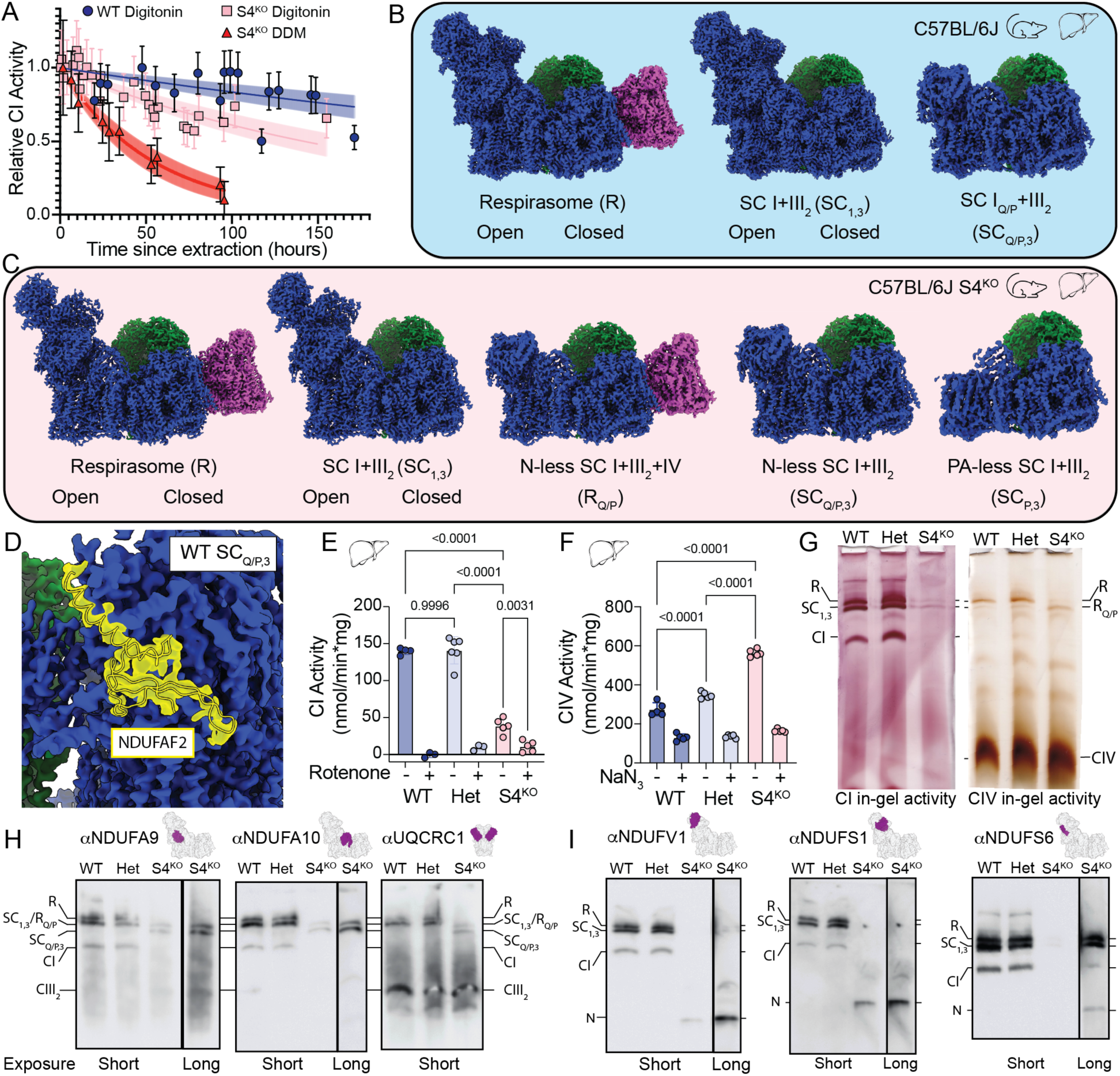
CI assembles on CIII_2_. (**A**) Time course of CI activity after detergent extraction from the WT and S4^KO^ mitochondrial membranes. (**B**) CryoEM density maps of respiratory supercomplexes isolated from WT mouse liver mitochondria colored with CI blue, CIII_2_ green and CIV magenta. (**C**) CryoEM density maps of respiratory supercomplexes isolated from S4^KO^ mouse liver mitochondria colored as in (**B**). (**D**) CryoEM density map of WT SC_Q/P,3_ showing the location of assembly factor NDUFAF2 colored as in (**B**) with transparent cryoEM density and model of NDUFAF2 shown in yellow. (**E**) CI NADH:decyl-ubiquinone activity from mouse liver mitochondria, n = 3-5, p-values from ordinary one-way ANOVA with multiple comparisons are shown. (**F**) Maximal CIV oxygen consumption driven by excess ascorbate, TMPD and cyt *c*, n = 4-5, p-values from ordinary one-way ANOVA with multiple comparisons are shown. (**G**) Blue Native (BN)-PAGE in-gel activity of digitonin extracted mouse liver mitochondrial complexes, left CI activity, right CIV activity. WT: NDUFS4^+/+^; Het: NDUFS4^+/-^; S4^KO^: NDUFS4^−/−^; supercomplex notations correspond to those shown in (**B**) and (**C**) with CI alone (CI) and CIV alone (CIV). (**H**) BN-PAGE western blots of digitonin extracted mouse liver mitochondrial complexes using antibodies against subunits present in SC_Q/P_, with primary antibody indicated at the top of the blot along with a structural representation of the subunit it targets highlighted in purple on the respiratory complex. Labels are as in (**G**) with CIII_2_ alone (CIII_2_). (**I**) BN-PAGE western blots of digitonin extracted mouse liver mitochondrial complexes using antibodies against N-module subunits present in fully assembled SCs. Primary antibody, subunit locations and labels are as in (**G**) with the N-module alone (N).

After initial cryoEM data processing we obtained five different SC structural classes for the WT samples: SC_1,3_ with CI in the open and closed states (*32*); R with CI in the open and closed states; and a minor class (3,000 particles) of SCs containing only the Q-module and the membrane arm proton-pumping (P) module of CI (Q/P intermediate) lacking the N-module (Figs. S1B, S3 and Tables S1, S2, S3). This subset contained both SC I_Q/P_+III_2_ (SC_Q/P,3_) and SC I_Q/P_+III_2_+IV (Q/P intermediate respirasome, R_Q/P_) particles but with too few total particles to separate (Fig. 1B, S3 and Table S3). The SC structures where CI was fully assembled and all subunits were present are equivalent to the recently published murine C-respirasome structures from both CD1 and C57BL/6J strains (*33*). CS-respirasomes with two copies of CIV were not observed, consistent with C57BL/6J mice lacking a functional SCAF1 subunit needed for interaction between CIII_2_ and CIV (*33*). Open and closed states of CI correspond mainly to the deactive and active states of the complex (*34*) and are present at roughly equal amounts in both SC_1,3_ and R consistent to what has been seen previously for isolated murine CI (*32*).

SCs formed between a CI subassembly lacking the N-module and CIII_2_ have been observed previously through faint bands on western blots (*14*) and as minor classes of SCs from ovine and bovine heart mitochondria (*35, 36*). However, due to the low number of particles it was unclear whether these were SCs containing CI assembly intermediates or degradation products. To determine this, we refined our WT SC_Q/P,3_ class to determine the presence or absence of CI assembly factor NDUFAF2. Despite only having 3,000 particles we were able to generate a 3.9 Å resolution focused map that unambiguously identified bound NDUFAF2 (Fig. 1D). Since NDUFAF2 is an assembly factor that is exchanged for subunit NDUFAF12 during final assembly of the peripheral arm (*37, 38*), its presence indicated that this SC subclass contains the CI_Q/P_ assembly intermediate (Fig. S1B), not a degradation product.

For the S4^KO^ samples we obtained maps where CI was fully assembled into either SC_1,3_ or R and maps with several classes of CI_Q/P_ assembled into either SC_Q/P,3_ or R_Q/P_ (Fig. 1C, S4 and Table S2, S3). The largest class of fully assembled CI SCs were missing NDUFS4, NDUFS6 and NDUFA12 and had NDUFAF2 bound (Fig. 1C and S5). This indicated that the majority of CI had not undergone the final step of assembly in which NDUFAF2 is replaced by NDUFA12. We also obtained fully assembled CI in both SC_1,3_ and R at a nominal resolution of 3.3 Å and 3.2 Å. In the WT, the ratio SC_1,3_:R was ∼2:1 while in the S4^KO^ this ratio was ∼1:1 indicating an increase in the relative abundance of CIV containing SCs (Figs. S3-S6 and Table S3). CI activity assays demonstrated a significant decrease in the CI activity of the mutant mitochondria, while maximal CIV activity was increased in the S4^KO^ livers and hearts relative to WT (Fig. 1E and S7A). This was not the case for CII activity which remained similar in WT, HET and S4^KO^ (Fig. S7B). These findings indicated increased CIV expression in the mutant, consistent with the observed higher ratio of R in the S4^KO^ (Table S3). In addition to structures with fully assembled CI, we identified two additional fully assemble CI classes, five different SC_Q/P,3_ subassemblies and one CI P-module-only (lacking the N- and Q-modules) bound to CIII_2_ (SC_P,3_, discussed further below).

Blue Native PAGE in-gel activity and western blots on digitonin extracted liver mitochondria samples confirmed that S4^KO^ CI is only found in SCs (Fig. 1G) (*14*). As a control, we performed a western blot using an antibody targeted against NDUFS4, which showed robust signal for respirasome, SC_1,3_ and CI alone in the WT and heterozygote (Het) samples but no signal in the S4^KO^ sample (Fig. S8A). Antibodies targeting CI subunits NDUFA9 and NDUFA10, which are present in fully assembled CI and CI_Q/P_, showed SC signal that correlated with R, SC_1,3_ and CI alone in the WT and Het samples (Fig. 1G, 1H, S8B, S8C). However, in the S4^KO^ lane we observed a band with lower molecular weight than the SC_1,3_ but larger than CI alone (Fig. 1G, 1H, S8B, S8C). A CIII_2_ specific antibody (αUQCRC1) showed bands consistent with those observed with the αNDUFA9 and αNDUFA10 antibodies indicating this band corresponds to a SC_Q/P,3_ (Fig. 1H, S8D). The CIV in gel activity assay also showed a band of similar size to SC_1,3_, consistent with R_Q/P_ (Fig. 1G). Further, we did not observed bands consistent with CI alone or CI_Q/P_ alone for the S4^KO^ sample (Fig. 1G, 1H, S8B, S8C), suggesting that CI_Q/P_ associates rapidly with CIII_2_ to form SC_Q/P,3_. When we used antibodies targeting the N-module subunits NDUFV1, NDUFS1 and NDUFS6 we observed clear bands for respirasome, SC_1,3_ and CI alone in the WT and Het samples (Fig. 1I, S8E, S8F, S8G). We observed only very faint bands consistent with fully assembled R and SC_1,3_ after long exposure with αNDUFV1 and αNDUFS6 antibodies in the S4^KO^ sample, but clear bands consistent with the N-module alone for all three N-module subunits consistent with CI instability or assembly defect (Fig. 1I, S8E, S8F, S8G). Overall, the in-gel activity and western blots confirmed that CI is only present in SCs in the S4^KO^ mitochondria and that the N-module alone, along with a SC_Q/P,3_ and R_Q/P_, accumulates in the S4^KO^, consistent with the final stages of CI assembly being delayed and occurring only after association with CIII_2_ (*14*).

### The S4^KO^ SC structures track the final steps of CI assembly

Classification of SC_Q/P_ and fully assembled SC particles from the S4^KO^ liver mitochondria resulted in a series of reconstructions that track the assembly and degradation of the CI peripheral arm (Fig. 2A, S5, S6). Initially, all R_Q/P_ and SC_Q/P,3_ particles were grouped together and 3D classified around the Q-module, revealing two major Q-module classes: one containing the minimal Q-module with NDUFAF2 bound, equivalent to the SC_Q/P,3_ class seen in WT (Fig. 1B, 1D), and the other containing the additional subunits NDUFA6 and NDUFAB1-α (SC_Q/P/A6,3_, Fig. 2, S6, S9). NDUFA6 and NDUFAB1-α are an LYRM/acyl-carrier protein pair that interact via the “flipped-out” acyl chain which is covalently bound to Ser44 of NDUFAB1-α and extends into the central cavity of NDUFA6 (*19, 39*) (Fig. 2F). NDUFA6 binds to the Q-module near the location of NDUFS4, positioning it above several key loops known to undergo conformational changes between the CI open and closed states (*34, 40*). The importance of NDUFA6 for CI activity has been demonstrated by several point mutations that significantly impact CI activity (*41*) including mimicking hypoxia by rescuing CI deficiency in S4^KO^ *C. elegans* (*31*). The fact that SC_Q/P/A6,3_ was not seen in the WT sample suggests that for WT CI the assembly step directly following formation of SC_Q/P,3_ is rate limiting, but attachment and full assembly of the CI N-module occurs rapidly thereafter. Thus, the absence of NDUFS4 in the KO introduces an additional rate limiting step, resulting in the buildup of two CI intermediates: SC_Q/P,3_ and SC_Q/P/A6,3_ (Fig. 2, S6).

**Figure 2.**
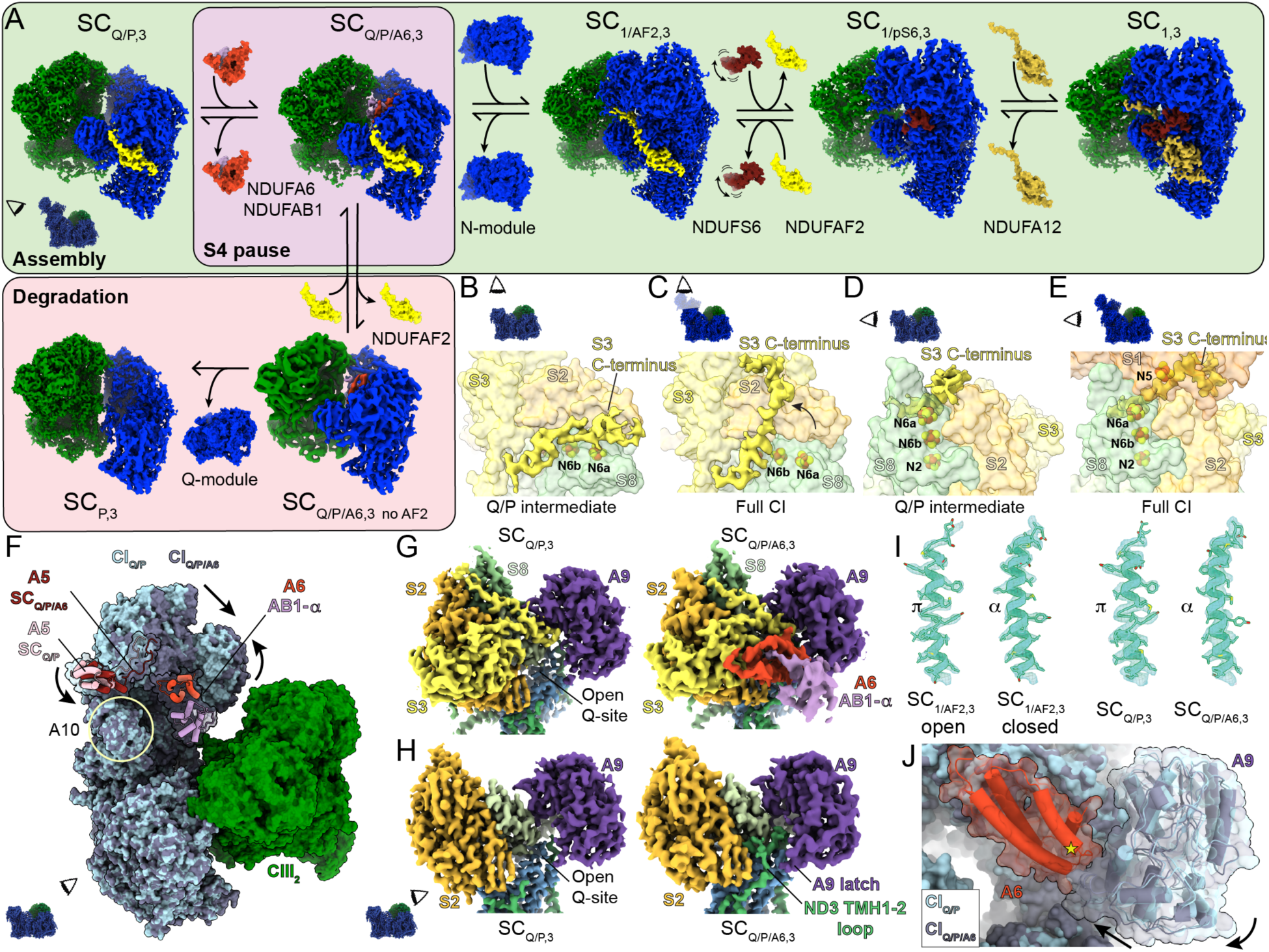
Assembly pathway and structural features of intermediates that accumulate in the S4^KO^. (**A**) CyoEM density maps of distinct SC assembly states generated from the S4^KO^ mouse liver data. The reconstructions are organized according to our model of CI assembly (light green box) and degradation (pink box). The additional Q/P assembly intermediate formed by binding of NDUFA6 (A6, red) and NDUFAB1-α (AB1-α, lilac) is highlighted with a purple background. (**B**) Matrix view of CI_Q/P_ intermediate (representative of both SC_Q/P,3_ and SC_Q/P/A6,3_) with the cryoEM map of the S3 C-terminus shown in yellow and the rest of NDUFS3 (S3), NDUFS8 (S8) and NDUFS2 (S2) shown in surface and labeled, CI FeS clusters are shown colored by atom (sulfur yellow and iron orange) and labeled. (**C**) Matrix view of full CI (representative of SC_1/AF2,3_, SC_1/pS6,3_ and SC_1,3_) with the N-module subunits hidden with S3 C-terminal density and subunits shown as in (**B**). (**D**) Back view of CI_Q/P_ intermediate (representative of both SC_Q/P,3_ and SC_Q/P/A6,3_) with NDUFAF2 hidden and S3 C-terminal density, FeS clusters and subunits shown as in (**B**). (**E**) Back view of full CI (representative of SC_1/AF2,3_, SC_1/pS6,3_ and SC_1,3_) with NDUFAF2/NDUFS6/NDUFA12 hidden with S3 C-terminal density and subunits shown as in (**B**) and NDUFS1 shown in light orange and label. (**F**) Overlay of SC_Q/P,3_ (CI_Q/P_ light blue, CIII_2_ green and NDUFA5 (A5) pink) and SC_Q/P/A6,3_ (CI_Q/P/A6_ dark blue, CIII_2_ green, A5 dark red, A6 red and AB1-α lilac). Structures were aligned by the MA and the difference in the position of the Q-module is indicated with arrows. Subunit NDUFA10 (A10) is circled and labeled. (**G**) CryoEM density map for Q-module subunits (NDUFS2 orange, NDUFS3 yellow, NDUFS8 sea green, NDUFA9 purple, NDUFA6 red, NUDFAB1-α lilac) and ND1 light blue and ND3 green of showing the position of SC_Q/P,3_ (left) and SC_Q/P/A6,3_ (right). (**H**) Similar to (**G**) with NDUFS3, NDUFS8, NDUFA6 and NDUFAB1-α removed to reveal the structured ND3 TMH1-2 loop and NDUFA9-latch in SC_Q/P/A6,3_, NDUFS7 density in pale green. Insets in (**A-H**) show viewing angle relative to the side view of the SCs with the matrix side up and CI blue and CIII_2_ green. (**I**) CryoEM density map and model for, from left to right, ND6 TMH3 from SC_1/AF2,3_ open, SC_1/AF2,3_ closed, SC_Q/P,3_ and SC_Q/P/A6,3_. The location and assignment of the ρε-bulge-α-helix transition is indicated. (**J**) Overlay of SC_Q/P,3_ (CI_Q/P_ light blue) and SC_Q/P/A6,3_ (CI_Q/P/A6_ dark blue and NDUFA6 red, NDUFAB1-α removed for clarity) showing the NDUFA9 conformational change induced by binding of NDUFA6). Star marks the equivalent position of the NDUFA6^G60D^ mutation in *C. elegans*.

In the SC_Q/P/A6,3_ class the density for subunit NDUFAF2 was significantly weaker than in the SC_Q/P,3_ class, prompting us to further classify using a mask around NDUFAF2 (Fig. S5B). This revealed a subset of ∼1.7 k particles that lack NDUFAF2, consistent with its disassociation (Fig. 2A, S6 and Table S3). Equivalent classification of the SC_Q/P,3_ class did not reveal a subset of particles lacking NDUFAF2, indicating that loss of NDUFAF2 occurs only after addition of NDUFA6 and NDUFAB1-α (Fig. 2A). This observation is consistent with the SC_Q/P/A6,3_ intermediate being a branching point between assembly and degradation of CI and suggests that the observed class SC_P,3_ is a further degradation product (Fig. 2A). Importantly, the CI P-module alone has not been identified as an assembly intermediate in studies of mammalian CI assembly (*9*) (Fig. S1). Two lines of evidence suggest that the putative CI degradation classes are not the result of our biochemical preparation but pre-exist within the mitochondrial membranes and thus represent native CI degradation from the stalled SC_Q/P/A6,3_ intermediate: 1) the loss of NDUFAF2 is state dependent, i.e., its absence from the SC_Q/P,3_, is only seen in the presence of the NDUFA6/NDUFAB1-α pair; and 2) the interface between CI and CIII_2_ within SCs can more easily be disrupted by harsh biochemical conditions than the interface between the CI MA and PA. Thus, it is unlikely conditions could remove the Q-module but leave CIII_2_ bound.

Next, we used masks around NDUFAF2 and the expected location of NDUFS6 to classify the intact SCs (S4^KO^ R plus S4^KO^ SC_1,3_; Fig. S5). This allowed us to separate three major classes: 1) NDUFAF2 bound SCs (S4^KO^ SC_1/AF2,3_), 2) partial NDUFS6 bound SCs (S4^KO^ SC_1/pS6,3_), and 3) NDUFS6 and NDUFA12 bound SCs (S4^KO^ SC_1,3_; Fig. 2A, S5, S6). These structures track the final stages of CI PA assembly. The NDUFAF2 bound state is an intermediate that accumulates in the absence of NDUFS4, as NDUFS4 helps dislodge NDUFAF2 by competing for binding at the interface of the N-module and Q-modules at NDFUS1 and NDUFA9 (Fig. S9A, S9B). As NDUFAF2 and NDUFS6 also clash (Fig. S9C), in the absence of NDUFS4, NDUFS6 binding alone must dislodge NDUFAF2. Additionally, as NDUFA12 and NDUFAF2 occupy the same binding site, NDUFAF2 must be dislodged before NDUFA12 can bind. This is complicated by the fact that the C-terminus of NDUFA12 binds underneath the N-terminal domain of NDUFS6 in the fully assembled complex (Fig. 2A). Confirming previous models (*26, 42*), this situation is resolved by the C-terminal domain of NDUFS6 binding first and dislodging NDUFAF2 while the N-terminal domain remains disordered allowing access for NDUFA12 to bind (S4^KO^ SC_1/pS6,3_; Fig. 2A and S9D). Only after NDUFA12 is bound does the N-terminal domain of NDUFS6 stably associate with the rest of the complex (S4^KO^ SC_1,3_; Fig. 2A). These results provide a series of structural snapshots along the pathway of CI PA assembly (Fig. 2A) and delineate the role of NDUFS4. NDUFS4 accelerates capture and association of the N-module, followed by destabilization of the assembly factor NDUFAF2 facilitating its exchange with NDUFS6 and NDUFA12.

### In CI_Q/P_ intermediates the C-terminus of NDUFS3 protects FeS cluster N6a

In all observed CI_Q/P_ assembly intermediates the C-terminal coil of core CI subunit NDUFS3 was found binding in a cleft formed on the solvent exposed surface between NDUFS8 and NDUFS2 (Fig. 2B, 2D). In fully assembled CI this cleft is occupied by the NDUFS1 subunit and the C-terminus of NDUFS3 binds across the surface of NDUFS2 (Fig. 2C, 2E). The location of the NDUFS3 C-terminus in SC_Q/P_ is directly above the FeS cluster N6a (4Fe[TY]1) (*43*), which is the first cluster in the Q module and would otherwise be nearly solvent exposed (Fig. 2D). This reveals a role for the NDUFS3 C-terminus in shielding N6a from oxidative damage during CI assembly. To complete the electron transport pathway and bring FeS cluster N5 (4F[75]H) within electron transfer distance to N6a, NDUFS1 must displace the NDUFS3 C-terminus from this cleft during attachment of the N-module (Fig. 2E). Consistent with this the density for the NDUFS3 C-terminus on SC_Q/P_ is partially disordered, suggesting multiple binding modes and overall low relative affinity for this site, which would facilitate its displacement during attachment of the N-module.

### The SC_Q/P/A6_ adopts a conformation consistent with reverse electron transport

The WT and S4^KO^ CI assembled with the N-module are found with CI in both open and closed states (Fig. 1B, 1C, S6, S10 and Table S3). The open state of CI has been shown to correspond to a catalytically inactive off-pathway state known as the deactive (D) state structurally characterized by disorder of loops forming the CoQ reduction site and a ρε-bulge in ND6 TMH3 (*34*) (Fig. S10). Although it is debated whether CI open states are also present on the catalytic cycle (*40, 44*), it has been clearly established that the D-state is open (*34, 45*). Deactive CI is not capable of catalyzing forward NADH:CoQ electron-transfer coupled proton-pumping (FET) or proton motive force (pmf) dissipating CoQH_2_:NAD^+^ reverse electron transfer (RET) (34). CI RET is a major source of reactive oxygen species (ROS) as O_2_ is a more energetically favorable electron acceptor than NAD^+^. Chemical modification or mutants that stabilize CI in the D-state can protect against ROS induced ischemia reperfusion injury by blocking CI RET (*46, 47*). When in the D-state CI must be reactivated by the addition of NADH to be competent for both FET and RET (*34, 48*). Structurally, the catalytic incompetency of the CI D-state can be understood through: the inability of CoQ to bind at it reduction site adjacent to FeS cluster N2 due to the disordered binding site loops; and a broken connection of hydrogen bonds and ordered water molecules between the CoQ reduction site and the hydrophilic axis of the membrane arm induced by the presence of the ND6 TMH3 ρε-bulge (*34, 49*). Previous structures of isolated CI from the S4^KO^ hearts and kidneys used chemical crosslinking to stabilize the complex and only observed CI in the closed state (*42*). However, in our rapid preparation we observe both open and closed states of CI in the SCs, indicating that CI lacking NDUFS4 deactivates (Fig. S4, S10). We confirmed the presence of the D-state in S4^KO^ heart mitochondria using an established N-ethyl maleimide (NEM) sensitivity assay (Fig. S11).

Both WT and S4^KO^ the CI_Q/P_ intermediates show structural characteristics expected for the D-state (Fig. 2F-J, 3A, S10, S12). When we pooled S4^KO^ SC_Q/P,3_ and R_Q/P_ particles (11,701 total) we were able to obtain a map at 3.3 Å resolution for CI that lacked density for the CoQ site loops (ND1 TMH5-6 loop, ND3 TMH1-2 loop, NDUFS2 β1-β2 loop, NDUFA9 latch, ND6 TMH3-4 loop and weak ND6 TMH4 density) and clearly showed a ρε-bulge in ND6 TMH3 (Fig. 2I). This indicates that the CI_Q/P_ intermediate is not catalytically competent as CoQ would not be able to bind adjacent to the N2 cluster and the hydrophilic axis is not engaged for proton-pumping (Fig. 3A, S10, S12). This is expected for an assembly intermediate that accumulates during WT CI assembly (Fig. 1B, S1B). However, this is not the case for the additional CI_Q/P/A6_ intermediate seen in the S4^KO^ SCs (Fig. 2A, S10, S12). Binding of the NDUFA6/NDUFAB1-α pair to SC_Q/P_ induces rotation of the Q-module relative to the MA and closing of the CoQ site (Fig. 2F-H, S12B, S12C). The conformational transition includes ordering of the ND1 TMH5-6 loop, the ND3 TMH1-2 loop, the NDUFS2 β1-β2 loop, the NDUFA9 latch and the conformational transition of ND6 THM3 into its α-helical form engaging the hydrophilic axis (Fig. 2F-I, 3B and S10). Further, as is commonly seen in the CI closed state, the rotation of the Q-module brings NDUFA5 into contact with NDUFA10 (*32*) (Fig. 2F, S12C). Thus, the conformation of the CI_Q/P/A6_ intermediate is consistent with the catalytically competent closed state. However, as the N-module is missing CI_Q/P/A6_ would only be able to catalyze RET, not FET. Thus, CI_Q/P/A6_ would only be capable of energy dissipation and its activity would be toxic to the cell. CI_Q/P/A6_ would establish a futile cycle that would dissipate energy, decrease the pmf and produce ROS (Fig. 3G) directly contributing to the pathophysiology of mitochondrial disease.

**Figure 3.**
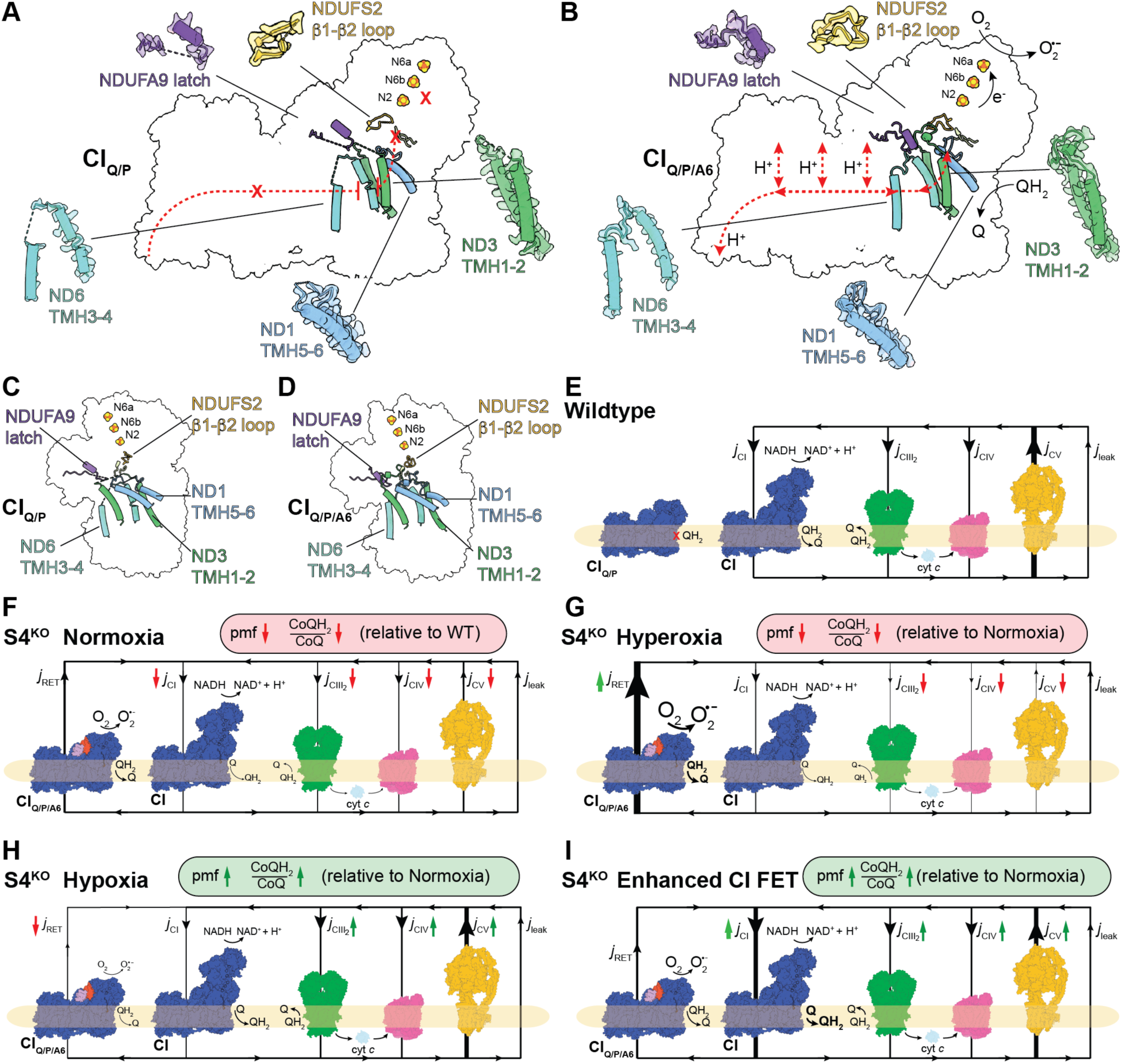
Pathophysiological consequences of active CI_Q/P/A6_ intermediate and implications for hypoxia rescue. (**A**) Side view of structural features of CI_Q/P_ intermediate consistent with the deactive state shown with matrix up. Catalytic incompetency indicated by red Xs, disordered regions of protein structure indicated by dashed lines. (**B**) Structural features of CI_Q/P/A6_ consistent with the active state. Catalytic competency indicated by arrows. (**C**) Similar to (**A**) shown from the back view. (**D**) Similar to (**B**) shown from the back view. (**E**) Schematic representation of mitochondrial H^+^ circuit. In the WT H^+^ flux 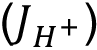 is rate limited by CV, allowing for the buildup of a large pmf that can be used to power ATP synthesis. (**F**) In S4^KO^, introduction of an oxygen dependent H^+^ leak (*j*_RET_) through CI_Q/P/A6_ under conditions of CI deficiency decreases the pmf, CoQH_2_/CoQ ratio and ATP synthesis relative to WT. (**G**) Hyperoxia in the S4^KO^ increases *j*_RET_ through CI_Q/P/A6_ further depressing the pmf, CoQH_2_/CoQ ratio and ATP synthesis relative to normoxia. (**H**) Hypoxia in the S4^KO^ decreases *j*_RET_ through CI_Q/P/A6_ improving the pmf, CoQH_2_/CoQ ratio and ATP synthesis relative to normoxia. (**I**) Enhanced CI FET in the S4^KO^ increases flux through the canonical chain (CIII_2_ and CIV) improving the pmf, CoQH_2_/CoQ ratio and ATP synthesis relative to normoxia.

## Discussion

### CIII_2_ acts as a platform for CI assembly

Two models for the assembly of CI and SCs have been debated in the field (*11, 13*). The assembly first plasticity model states that the complexes must fully assemble prior to association into SCs (*9*), whereas the cooperative assembly model proposes that partially assembled complexes can associate before they are fully assembled (*10–13*) (Fig. S1B). Our structures strongly support the cooperative assembly model (Fig. 1B, 1C). These structures showed partially assembled CI bound to CIII_2_ or CIII_2_ and CIV, indicating that CI does not need to finish assembly prior to SC formation. In addition, since NDUFAF2 is seen bound to SC_Q/P,3_ in the WT sample (Fig. 1B, 1D) our data demonstrates that although the assembly process is slowed in the S4^KO^, resulting in the accumulation of several additional intermediates, cooperative assembly occurs in the WT. This is consistent with studies of induced CIII_2_ deficiencies resulting in concomitant CI deficiency and the accumulation of a CI_Q/P_ intermediate (*11, 28*). Further, mutations in NDUFA6 have also been shown to accumulate SC_Q/P,3_ and R_Q/P_ intermediates with NDUFAF2 bound (*50*). These observations along with our structural data indicate that the attachment of the N-module to CI_Q/P_ is the rate limiting step in WT mammalian CI assembly, leading to a steady state buildup of this intermediate. Since induced CIII_2_ deficiency stalls CI assembly at CI_Q/P_ (*11, 51*), these data indicate that CIII_2_ acts prior to NDUFS4 and NDUFA6, likely promoting their association, which is rapidly followed by the full N-module. Cooperative assembly has also been observed in structures of mouse SC III_2_+IV (*52*), suggesting that this is a general approach for mammalian respiratory complex assembly.

### The toxic CI_Q/P/A6_ intermediate explains mitochondrial disease rescue by hypoxia

The Mootha and Zapol labs demonstrated that chronic hypoxia improves survival, body weight, body temperature, behavior, neuropathology and disease biomarkers in the S4^KO^ mice (*29, 30*). Further they showed that hypoxia treatment can reverse neurodegeneration in these mice (*30*), whereas hyperoxia exacerbates disease (*29*). More recently, Meisel *et al.* demonstrated that hypoxic rescue of S4^KO^ is conserved in *C. elegans* and that mutations in subunits NDUFA5 and NDUFA6 phenocopy hypoxia rescue (*31*). Our structure of the SC_Q/P/A6,3_ intermediate structurally competent for O_2_-dependent RET accounts for these observations. Due to the coupled nature of the reaction catalyzed by CI, oxidation of CoQH_2_ by SC_Q/P/A6,3_ would depend on proton transport across in the IMM. The exact mechanism of electron transfer coupled proton pumping is still debated (*49, 53–55*) but it is well established that both are needed for CI FET and RET (*48*). As CI_Q/P/A6_ could only perform RET, energy from the pmf will be dissipated (Fig. 3G). WT CI can catalyze the thermodynamically unfavorable reduction of NAD^+^ by CoQH_2_ using energy from the pmf (48). However, if O_2_ is present, CI can catalyze the pmf-powered reduction of O_2_ by CoQH_2_ generating superoxide (*56*). Given that both dissipation of the pmf and the electron transfer from CoQH_2_ to O_2_ are energetically favorable, RET by CI can be a major source of ROS, leading to tissue damage in ischemia reperfusion injury or metabolic dysfunction and chronic disease (*57, 58*). In the case of RET by CI_Q/P/A6_, electron transfer to NAD^+^ is not possible as the NADH binding site is absent (Fig. 3B). However, an electron acceptor is still needed to support toxic RET flux (*j*_RET_) through the intermediate (Fig. 3G). CI_Q/P/A6_ has three of the seven FeS clusters (N2, N6b and N6a) that form the electron transport pathway between NADH and CoQ (Fig. 3B). Each of these FeS clusters could accept one electron, but in the NADH reduced enzyme only N6a and N2 are seen reduced simultaneously (*59, 60*). This indicates that the three Q-module clusters can only accept two electrons at a time, i.e. oxidize only a single CoQH_2_ at a time. This single CoQH_2_ oxidation would be coupled to pmf dissipation, but subsequent CoQH_2_ oxidations needed to generate significant *j*_RET_ would require oxidation of the FeS clusters by O_2_. This reaction would produce ROS while oxidizing the Q-module FeS clusters allowing for further rounds of pmf dissipation coupled CoQH_2_ oxidation (Fig. 3G). Thus, sustained CoQH_2_ oxidation and energy dissipation by SC_Q/P/A6,3_ would be O_2_ dependent.

The flow of protons across the IMM that is catalyzed by the oxidative phosphorylation complexes form a circuit that can be understood analogously to a simple electrical circuit (*61, 62*) (Fig. 3F). Under normal conditions H^+^ current 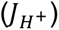 is rate limited by H^+^ flux into the mitochondrial matrix through the ATP synthase (*j*_CV_) and H^+^ leak (*j*_leak_) allowing for the proton-pumping complexes (CI, CIII_2_ and CIV) to buildup of a large pmf (Fig. 3F). Introduction of additional leak through CI_Q/P/A6_ (*j*_RET_) would lower the pmf by increasing H^+^ conductance across the membrane (Fig. 3G). A lower pmf would normally increase 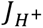 as reactions catalyzed by CI, CIII_2_ and CIV would face lower resistance from the membrane electrochemical potential. However, under conditions of CI deficiency, the four-fold decrease in CI (Fig. 1E) would greatly reduce overall CI flux (*j*_CI_). Further, the CoQH_2_:O_2_ oxidoreduction catalyzed by CI_Q/P/A6_ introduces a competing pathway for CoQH_2_ oxidation, slowing flux through CIII_2_ (*j*_CIII2_) and CIV (*j*_CIV_), diminishing the CoQH_2_/CoQ ratio and exacerbating the CI deficiency (Fig. 3G). Diminished pmf and CoQH_2_/CoQ ratio have been observed S4^KO^ *C. elegans* (*31*). In this model, the energy dissipating CI_Q/P/A6_ reaction is rate limited by O_2_ availability explaining how hyperoxia and hypoxia modulate pathophysiology (*29, 30*). Hyperoxia would promote and hypoxia would block *j*_RET_ at CI_Q/P/A6_ (Fig. 3H and 3I). Thus, CI_Q/P/A6_ *j*_RET_ can contribute to disease in two ways: 1) by producing of ROS and 2) by establishing a futile respiratory cycle that dissipates pmf, leading to a lower CoQH_2_/CoQ ratio and a lower steady-state rate of ATP synthesis (Fig. 3G). Importantly, Meisel *et al.* ruled out decreased mitochondrial ROS toxicity underlying the rescue by hypoxia in *C. elegans* (*31*), suggesting that the true pathological defect rescued by hypoxia is the aberrant dissipation of the pmf by CI_Q/P/A6_ (Fig. 3G and 3I).

In the context of the RET-competent structure of SC_Q/P/A6,3_, the hypoxia mimicking NDUFA5^R126Q^ and NDUFA6^G60D^ mutants in *C. elegans* are understood not through their impact on fully assembled CI, but through their ability to impede CI_Q/P/A6_ adopting a RET competent state, i.e., ordering the Q-site loops and inducing the ρε-bulge-to-α-helix transition in ND6 THM3 (Fig. 2F-I, S10, S12). The conformational change between open Q-site CI_Q/P_ and closed Q-site CI_Q/P/A6_ requires a rotation of the Q-module relative to the P-module that results in protein-protein interactions between NDUFA5 and NDUFA10 (Fig. 2F, S12). The residue equivalent to *C. elegans* NDUFA5^Arg126^ on mouse NUDFA5 makes several charge-charge interactions with acidic residues on NDUFA10 in the closed state (Fig. S12C). Thus, mutation of NDUFA5^Arg126^ to the neutral glutamine would destabilize the closed state of the CI_Q/P/A6_ intermediate decreasing the likelihood of it adopting a RET competent state, resulting in the rescue phenotype. As the binding of NDUFA6/NDUFAB1-α to CI_Q/P_ induces closing of the intermediate into a RET competent state, the effect of the NDUFA6^G60D^ mutation can be understood as decreasing the ability of NDUFA6 to induce this conformational change (Fig. 3A and 3B). Disordering of the CI Q-site loops have been observed in structures of the *Yarrowia lipolytica* NDUFA6(NUYM)^F89A^ mutant (*63*), which is structurally adjacent to the position of the G60D mutant in *C. elegans* (Fig. S12D, S12E, S12F). Consistent with this, NDUFA6^G60D^ sensitizes CI to the Q-site inhibitor rotenone highlighting its structural impact on the CoQ binding site (*31*). Importantly, rotenone is a competitive inhibitor of CoQ and thus increased rotenone sensitivity could be induced by either increasing affinity for the inhibitor itself or by decreasing the affinity for the competing substrate, CoQ. Another mechanism that could overcome *j*_RET_ through CI_Q/P/A6_ would be to enhance FET at CI (Fig. 3J). Increasing the rate of CI FET, would increase the steady-state level of the pmf and CoQH_2_/CoQ as was observed for the NDUFS2^R290K^/ NDUFA6^G60D^ double mutant in *C. elegans* (*31*).

## Conclusions

The characterization of CI deficiencies has relied on using intact CI structures to interpret molecular, cellular and organismal physiological data. By structurally characterizing SCs from the S4^KO^ mouse model of Leigh syndrome we show: first, that CI is co-operatively assembled by CIII_2_; second, that in the absence of NDUFS4 aberrant assembly intermediates accumulate; and third, that it is these intermediates, rather than merely a lack of fully assembled CI, that are pathophysiological. This reveals a new mechanism underlying mitochondrial disease in which aberrant CI assembly intermediates dissipate the pmf via an O_2_-dependent reaction that leads to a lower CoQH_2_/CoQ ratio and a lower steady-state rate of ATP synthesis. Going forward careful characterization of the CI assembly state will be needed to fully understand the pathophysiology of CI deficiencies in mitochondrial disease.

## Acknowledgments

Some of this work was performed at the Stanford-SLAC Cryo-EM Center (S^2^C^2^), which is supported by the National Institute of General Medical Sciences (1R24GM154186). The content is solely the responsibility of the authors and does not necessarily represent the official views of the National Institutes of Health. The authors would also like to thank the following S^2^C^2^ personnel for their invaluable support and assistance: Dr. Ian Fries and Dr. Patrick Mitchell.

## Author contributions

Conceptualization: GC, JAL

Methodology: MGA, JAL

Investigation: MGA, ABT, HCT, CBM, AP, JAL

Visualization: MGA, AP, JAL

Funding acquisition: MGA, GC, JAL

Project administration: GC, JAL

Supervision: GC, JAL

Writing – Original Draft: MGA, JAL

Writing – review & editing: MGA, CBM, GC, JAL

## Data and materials availability: Data and materials availability

Single particle cryogenic electron micrograph movies and motion corrected micrographs for WT and S4^KO^ are available on the Electron Microscopy Public Image Archive, accession codes: EMPIAR-12845 and EMPIAR-12846, respectively. The maps and models for WT and S4^KO^ SCs are available on the Electron Microscopy Database (EMDB) and the Protein Data Bank (PDB) with accession codes: EMDB-71223, PDB-9P30; EMDB-71222, PDB-9P2Z; EMDB-71220, PDB-9P2X; EMDB-71221, PDB-9P2Y; EMDB-71224, PDB-9P31; EMDB-71219, PDB-9P2W; EMDB-71218, PDB-9P2V; EMDB-71225, PDB-9P32; EMDB-71122, PDB-9PIL; EMDB-71214, PDB-9P2S; EMDB-71216, PDB-9P2T.

## Supplementary Materials

### Material and Methods

#### Animals and genotype analysis

All animal protocols were approved by the Institutional Animal Care Use Committee at the University of California, Davis and were also in accordance with the NIH guidelines for the Care and Use of Laboratory Animals. The NDUFS4^+/−^ mouse strain on a C57BL/6J background was provided by the Jackson Laboratory (Bar Harbor, ME) and purchased for in-house breeding. To produce constitutive NDUFS4 knockout mice, mice heterozygous for the NDUFS4 knockout (NDUFS4*^+/−^*) were bred together. Wild-type littermates (NDUFS4*^+/+^*) were used as controls and heterozygotes (NDUFS4*^+/−^*) were also assessed in parallel. Male and female mice were housed in polycarbonate cages starting at 21 days of age on a 12-hour light/dark cycle. Body weights were monitored weekly. Mice were provided DI water and a standard rodent chow *ad libitum* (Teklad 2018, Inotivco). Mice were provided with Mouse Igloos (Bio-Serv) to provide additional shelter. Mice were euthanized by carbon dioxide overdose at 43 days of age and intra-cardiac perfusion performed with 5 mL of phosphate buffered saline before removal of target organs. We prepared genomic DNA from ear clip samples collected from the mice between 12-20 days of age by using a HotShot DNA extraction (using 0.5M NaOH and Tris HCl pH 7.0 solutions), and PCR-based genotyping was completed using the published primer sets (*24*) for wild type and knockout alleles (Integrated DNA Technologies).

#### Mitochondrial purification

Liver tissues were excised from wild-type and NDUFS4 knockout C57BL/6J mice and rinsed in PBS before blotting them dry. The tissues were then flash frozen in liquid nitrogen and stored at −70 °C. All steps were performed at 4°C with pre-chilled materials. Mitochondria were isolated from the knockout and wildtype tissues as described previously (*64*). Briefly mouse livers were thawed at 4 °C before being minced into 1 mm pieces and homogenized in 10 mL buffer AT (0.075 M sucrose, 0.225 M sorbitol, 1 mM EGTA, 0.1% fatty acid-free bovine serum albumin (BSA), and 10 mM Tris-HCl, pH 7.4, supplemented with SIGMAFAST Protease Inhibitor Cocktail Tablets) per gram of tissue using a Potter-Elvehjem homogenizer fitted with a Teflon pestle for 50 strokes. The homogenate was spun down at 1,000 g for 5 min. The supernatant from that spin was transferred to eight 1.5 mL Eppendorf microcentrifuge tubes and spun at 15,000 g for 2 min. The supernatant was removed, leaving only the brown mitochondrial pellet. Two mitochondrial pellets were combined dividing the number of tubes in half and were resuspended in 1.5 mL of medium AT using a pipette. The combined pellets were spun down at 15,000 g for 2 min. The process was repeated until there was only 1 pellet in one tube. The supernatant was removed from the final pellet and the pellet was considered crude mitochondria and stored at −70 °C.

#### Mitochondrial membrane wash

All steps were performed at 4°C with pre-chilled materials. Unfrozen mitochondria from the mitochondrial purification step were homogenized in 10 mL MilliQ water per gram of mitochondria using a Dounce glass homogenizer for 100 strokes. Potassium chloride was added to final concentration of 0.15 M and sample was homogenized again for 100 strokes. The homogenate was centrifuged at 43,667 g for 50 min. The pellet was resuspended in 18 mL Buffer M (20 mM Tris pH 7.4, 50 mM NaCl, 1 mM EDTA, 10% (v/v) Glycerol, 2 mM DTT, 0.002% PMSF) per gram of mitochondria starting material for and homogenized again for 100 strokes. The homogenate was centrifuged at 28,302 g for 50 min. The pellet was resuspended in 3 mL Buffer M (20 mM Tris pH 7.4, 50 mM NaCl, 1 mM EDTA, 10% glycerol, 2 mM DTT, 0.002% PMSF) per gram of mitochondrial starting material and homogenized again for 100 strokes. Protein concentration of the final membranes was calculated using a Pierce BCA assay kit and was diluted to final concentration of 10 mg/ml in 30% (v/v) glycerol for storage at −70 °C.

#### Supercomplex purification

Mouse liver washed mitochondrial membranes were thawed on ice. Supercomplex were isolated by tumbling for 45 min at 4 °C with digitonin at a 4:1 (w/w) ratio and 1% (w/v) concentration in Buffer MX (30 mM HEPES, 150 mM potassium acetate, 10% v/v glycerol, 1 mM EDTA and 0.002% PMSF). The sample was centrifuged at 15,973 *g* for 45 min at 4 °C and the supernatant was kept. The supernatant was concentrated using (100 kDa cutoff) centrifugal concentrators to a final volume of 500 µl (knockout) and 250 µl (wildtype) to inject onto the size exclusion chromatography (SEC) Superose 6 increase 10/300 GL column pre equilibrated with SEC buffer (30 mM HEPESpH 7.8, 150 mM potassium acetate, 1 mM EDTA, 0.005% (w/v) GDN). Fractions were run on a 3-12% BN-page and subjected to CI-in-gel activity assay.

Fractions showing CI activity were pooled for both the knockout and wildtype samples. The pooled fractions were concentrated using (100 k Da cutoff) centrifugal concentrators and protein concentration was measured using a Pierce BCA assay kit and diluted to ∼5 mg/ml in 0.05 % digitonin in Buffer MX to freeze cryoEM grids.

#### Cryo-EM grid preparation and data collection

For the WT, data was collected from two different cryoEM grids. Three microliters of 6 mg/mL fractions from Superose 6 increase 10/300 GL column were applied onto a C-Flat 1.2/1.3 20 nm carbon on 300 mesh gold grid, glow discharged at 30 mA for 20 seconds. The grids were incubated with sample for 10 seconds pre-blotting at 20 °C and 90% humidity. One grid was blotted for 7 seconds and the other for 8 seconds before plunge-freezing into liquid ethane. A total of 16,310 movies were collected using EPU on a 300 kV Titan Krios with a pixel size of 0.86 Å/pixel. A dose of 50.5 electrons/Å^2^ with a 1.68 s exposure time was fractionated into 40 frames for each movie.

For the S4^KO^, data was collected from three different cryoEM grids. Three microliters of 6 mg/ml fractions from Superose 6 increase 10/300 GL column were applied onto a C-Flat 1.2/1.3 20 nm carbon on 300 mesh gold grid, glow discharged at 30 mA for 20 seconds. The grids were incubated with sample for 10 seconds pre-blotting at 20 °C and 90% humidity. Two grids were blotted for 10 seconds and the other for 6 seconds before plunge-freezing into liquid ethane. A total of 17,190 movies were collected using EPU on a 300 kV Titan Krios with a pixel size of 0.86 Å/pixel. A dose of 49.78 electrons/Å^2^ with a 1.63 s exposure time was fractionated into 40 frames for each movie.

#### Cryo-EM image pre-processing for knockout and wildtype sample

The raw movies were motion-corrected using MotionCor2 and per-micrograph contrast transfer function (ctf) estimation was calculated using the CTFFIND4.1 in Relion 4.1.0. Using Warp (*65*), micrographs were curated to remove 556 in the WT and 958 in the S4^KO^ datasets. Particles were picked using a trained model. The initial 1,215,504 WT and 1,424,255 S4^KO^ picked particles were extracted in Warp with 600 pixel^2^ boxes and imported into cryoSPARC v4.4.1. Iterative 2D classification, 3D ab initio reconstruction, and 3D refinement were performed initially in CryoSPARC.

In the WT sample, 117,010 good particles were obtained after the final round of 2D classification. 3D *ab initio* and 3D classification resulted in 61,454 particles corresponing to SC I+III_2_ and 31,935 to the respirasome. Homogeneous refinement and non-uniform refinement of each of the classes resulted in reference map of 3.5 Å. The particle set was then transferred back into Relion 4.1.0 for global search, CTF refinement, Bayesian polishing and local searches resulting in a final map of 3.0 Å for SC I+III_2_ and 3.1 Å for the respirasome. These maps were used for initial model building in coot and refinement in phenix.

In the S4^KO^ sample, 77,586 good particles were obtained after the final round of 2D classification. 3D *ab initio* and 3D classification resulted in 88,714 particles in 18 classes that corresponded to 14,547 SC I+III_2_, 18,452 to the respirasome, 8,007 to SC I_Q/P_+III_2_ and 8,037 to R_Q/P_. Lastly, heterogenous refinement of particles missing the N-module yielded a class of 3,000 particles without peripheral arm. Homogeneous refinement and non-uniform refinement of each of the classes resulted in reference map of 3.4 Å. The particle set was then transferred back into Relion 4.1.0 for global search, CTF refinement, Bayesian polishing and local searches resulting in a final map of 3.3 Å for SC I+III_2,_ 3.5 Å for the respirasome, 3.8 Å for N-less SC I+III_2_ and 3.9 Å for R_Q/P_. These maps were used for initial model building in coot and refinement in phenix.

#### Model building and refinement

Model building was performed in Coot and refinements in Phenix 1.21. For the WT, CI structure from murine (6ZR2) and SC III_2_+IV structure from murine (7O3C) were docked into our structures. For the S4^KO^, CI structure from murine (8CA5) and SC III_2_+IV structure from murine (7O3C) were docked into our structures. The models, were manually inspected, adjusted, and rebuilt where necessary to generate our model.

#### Blue native PAGE

Mouse liver mitochondrial membranes were solubilized using 1% digitonin for 45 min and centrifuged at 16,130 g for 30 min. The supernatant was concentrated, and 40 µg of total protein were loaded on Bio-Rad 4-15% Mini-PROTEAN TGX Precast Gels. The gel was run for 30 minutes at 150 V in buffer containing 0.02% Coomassie Brilliant Blue G. After, the gel was run for 1 hour and 30 minutes at 200 V in a buffer containing 1/10^th^ of the Coomassie Brilliant Blue G buffer. In-gel complex I activity assays were performed using 150 µM NADH and 1.5 mg/mL Nitrotetrazolium Blue chloride (NTB). In-gel complex IV activity assays were performed using 10 mM phosphate buffer pH 7.4, 50 mM NaCl, 0.5 mg/mL 3,3’-diaminobenzidine (DAB) and 80 µM cytochrome *c*.

#### Western blotting

Protein complexes were separated using the blue native PAGE methods described above. The proteins were transferred to polyvinylidene difluoride (PVDF) membranes using a Transblot Turbo Transfer System (Bio-Rad). The PVDF membrane was blocked in 5% (w/v) nonfat dry milk in tris-buffered saline (TBS) overnight. The membranes were washed twice for 10 min each in 0.05% Tween 20 in TBS (TBST). Following the washes, the membranes were incubated in the appropriate primary antibody dissolved in 5% (w/v) nonfat dry milk in TBST for 2 hours at room temperature. The membranes were washed four times for 10 min each in TBST. After the washes, the membranes were incubated for 1 hour at room temperature with the appropriate HRP-conjugated secondary antibody dissolved in TBST. Then, the membranes were washed four times for 10 min each in TBST. A final wash was performed in TBS to remove Tween 20 from the membrane surface. Immunoreactivity was detected by a Prometheus Protein Biology Products ProSignal Femto kit (Genese Scientific) and analyzed by the Lumenescent Image Analyzer (Image Quant LAS400). Protein immunodetection was performed using the primary antibodies: anti-NDUFS6 (ab195807, Abcam), anti-NDUFS4 (ab139178, Abcam), anti-NDUFA10 (PA5-22348, Invotrogen), anti-NDUFA9 (459100, Abcam), anti-CORE Protein I (ab110252, Abcam), anti-NDUFS1(PA5-22309, Invitrogen). The secondary antibodies used were: goat anti-rabbit IgG (ab6721), and goat anti-mouse (AP181P, EMD Millipore).

#### Complex I Activity

Complex I activity was measured from wildtype and knockout murine liver mitochondrial membranes in reaction buffer (20 mM HEPES pH 7.4, 50 mM NaCl, 10% (w/v) glycerol, 0.1% (w/v) CHAPS, 0.25 mg/mL 4:1 Asolectin:Cardiolipin, 1 mg/mL BSA, 100 µM Decylubiquinone) at 200 mg by measuring NADH oxidation at 340 nm in 4.5 mL cuvettes at room temperature using the Cary 60 UV-Vis (Agilent). Mitochondrial membranes were mixed with reaction buffer using a stir bar to a final volume of 1.5 mL. The reaction was initiated by the addition of 150 µM NADH and Rotenone (1 µM) was used to inhibit CI and show specificity. Measurements of the initial rates were done in triplicates, averaged and normalized. A CI assay as described above was used to measure the stability of CI over time after extraction in either Digitonin or DDM.

#### Complex IV Activity

Complex IV activity was measured from wildtype, heterozygote, and knockout murine liver and heart mitochondrial membranes in reaction buffer (20 mM HEPES pH 7.4, 50 mM NaCl, 10% (w/v) glycerol, 0.1% (w/v) CHAPS, 0.25 mg/mL 4:1 Asolectin:Cardiolipin, 1 mg/mL BSA) at 200 mg by measuring oxygen consumption using an Oxygraph+ (Hansatech Instruments Ltd). The reaction buffer was added to the Oxygraph+ chamber with cytochrome *c* (100 µM) and the mitochondrial membranes and was constantly mixed using a stir bar. The reaction was initiated by the addition of TMPD (300 mM) and ascorbate (3 mM). Sodium Azide (1 mM) was used to inhibit complex IV. Measurements of the oxygen concentration were done in a minimum of triplicates, averaged and normalized.

#### Complex II Activity

Complex II activity was measured from wildtype, heterozygote, and knockout murine liver mitochondrial membranes in reaction buffer (50 mM HEPES pH 8.0, 0.1 mM EDTA, 1 mg/mL BSA, 0.25 mg/mL 4:1 Asolectin:Cardiolipin, 4 µM KCN, 1 µM Rotenone, 2 µM Antimycin, 100 µM Decylubiquinone) at 200 mg by measuring DCPIP reduction at 600 nm using the Cary 60 UV-Vis (Agilent). Measurements were made in 4.5 mL cuvettes at room temperature at a final volume of 1.5 mL with constant stirring using a stir bar. Membranes were added to the buffer and were allowed to equilibrate, DCPIP (100 µM) was added, and the reaction was initiated by the addition of succinate (100 µM). Oxaloacetate (200 µM) was used to inhibit complex II to show specificity. Measurements of the initial rates were done in a minimum of triplicates, averaged and normalized.

#### NEM Assay

The NEM assay was performed in 96 well plates for wildtype and knockout murine liver and heart mitochondrial membranes in reaction buffer (20 mM HEPES pH 7.4, 50 mM NaCl, 10% (w/v) glycerol, 0.1% (w/v) CHAPS, 0.25 mg/mL 4:1 Asolectin:Cardiolipin, 1 mg/mL BSA, 100 µM Decylubiquinone) at 300 µg for the knockout and 30 µg for the wildtype by measuring NADH reduction at 340 nm. Mitochondrial membranes from wildtype and knockout hearts and livers were incubated at 37 °C for 30 (wildtype) and 15 (knockout) minutes or left as is. 5 µM pre NADH or an equivalent amount of buffer was added to the corresponding sample and mixed by pipetting. 30 seconds after the addition of pre NADH or buffer 2 mM NEM or water was added to the corresponding well and mixed by pipetting. The plates were covered and incubated at room temperature for 20 minutes. The reaction was started by the addition of 200 µM NADH and NADH oxidation was measured at 340 nm using a Molecular Devices (San Jose, CA) Spectramax M2 spectrophotometer. Measurements of the initial rates were done in triplicates, averaged and normalized.

### Supplementary Figures

**Figure S1.**
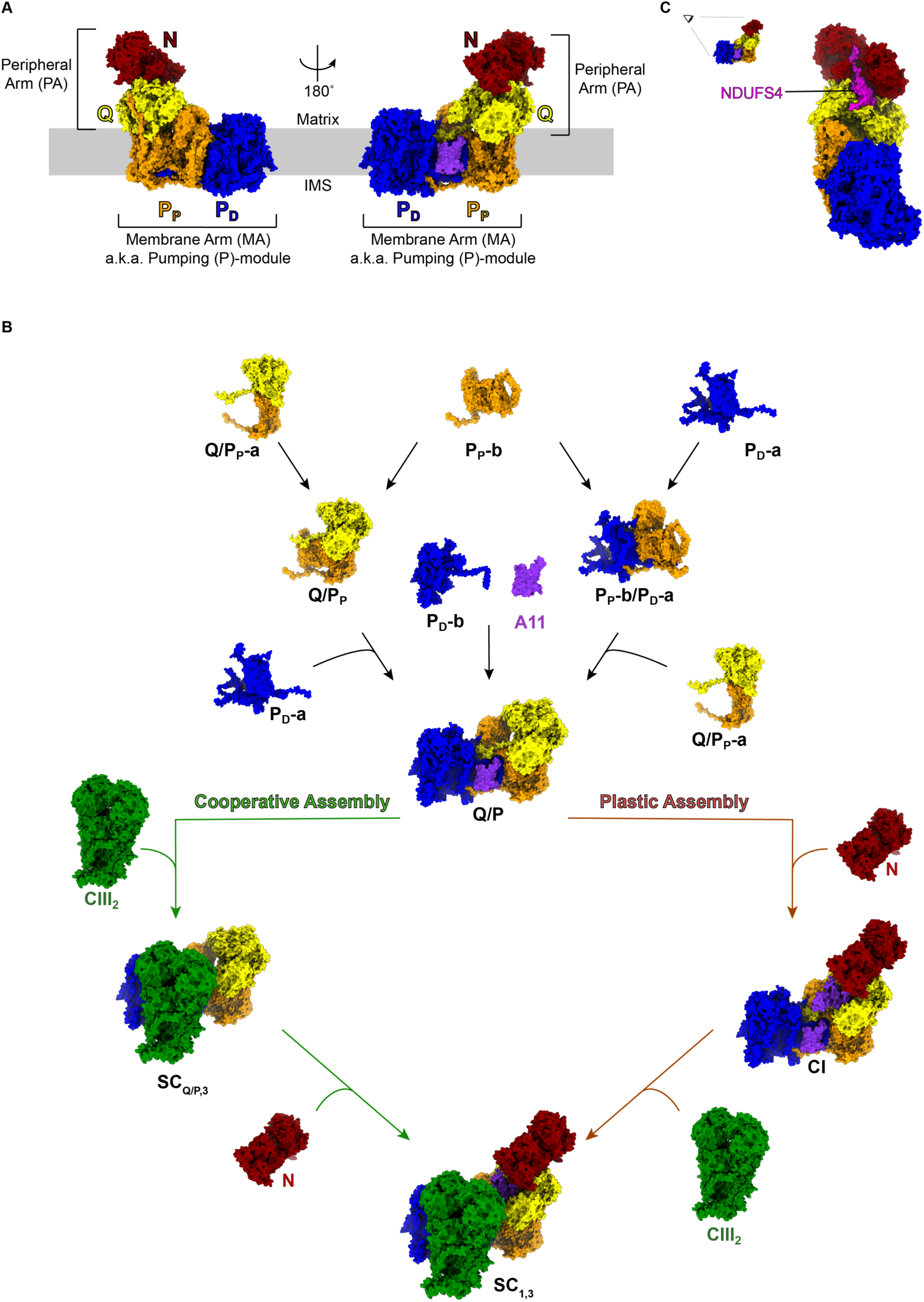
Modular nature of CI structure. (**A**) Mouse WT CI structure colored by module with the different arms of the complex indicated. (**B**) Simplified modular assembly pathway for CI comparing the cooperative assembly model vs. assembly first plasticity model. (**C**) Location of subunit NDUFS4.

**Figure S2.**
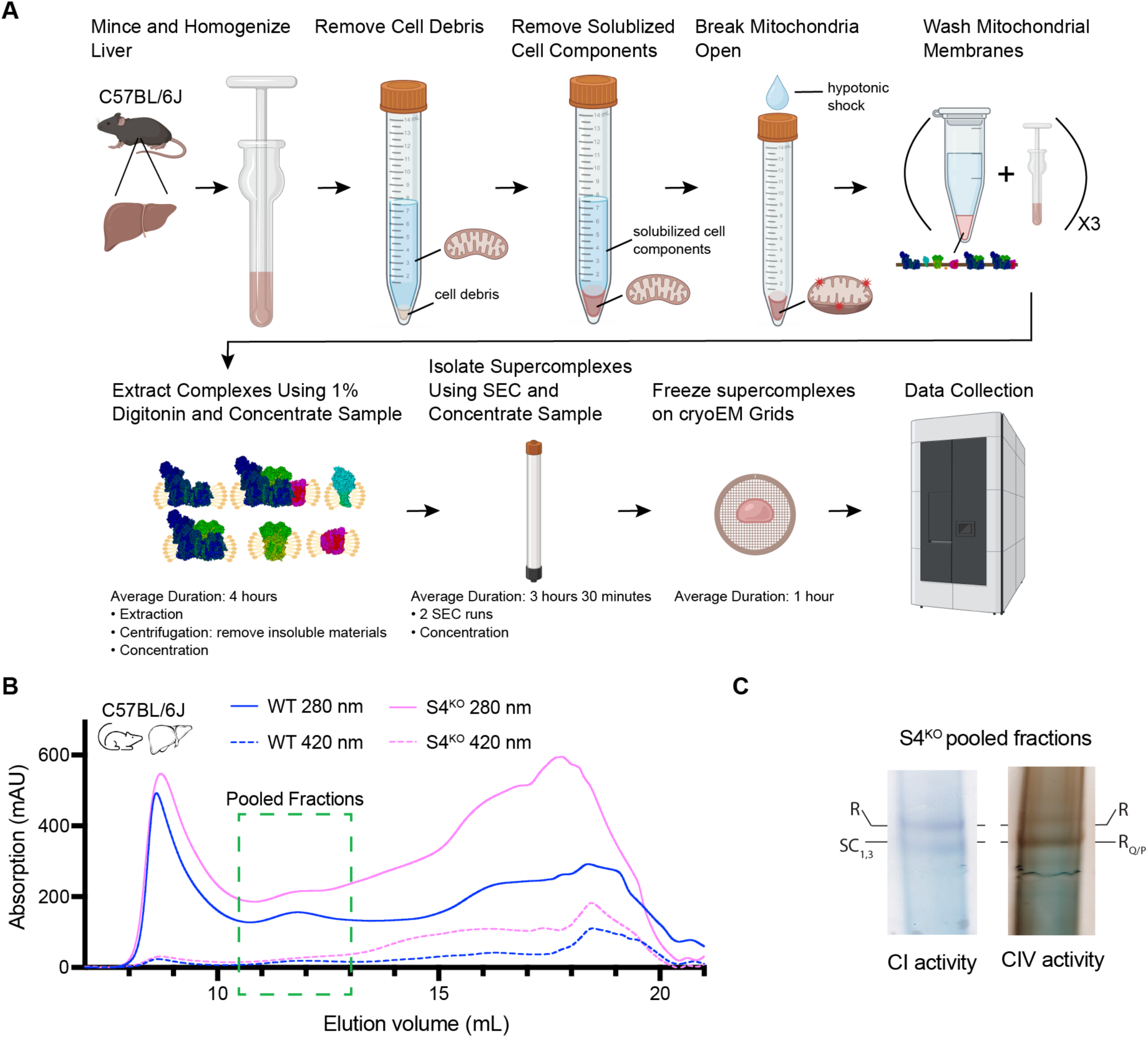
Biochemical preparation of SC samples. (**A**) Schematic showing the steps from tissue homogenization to sample freezing on grids. (**B**) Superose 6 Increase 10/300 size exclusion column (SEC) chromatogram of 1% digitonin (w/v) extracted mitochondrial membranes from liver. WT: NDUFS4*^+/+^* and S4^KO^: NDUFS4*^−/−^*. The green dotted box indicates the fractions that were pooled and concentrated for grid freezing. (**C**) Blue-native PAGE (BN-PAGE) CI (left) and CIV (right) in-gel activity assays of S4^KO^ pooled fractions from (**B**). Labels: R: Respirasome; SC_1,3_: Supercomplex I+III_2_; R_Q/P_: Respirasome containing CI Q/P intermediate.

**Figure S3.**
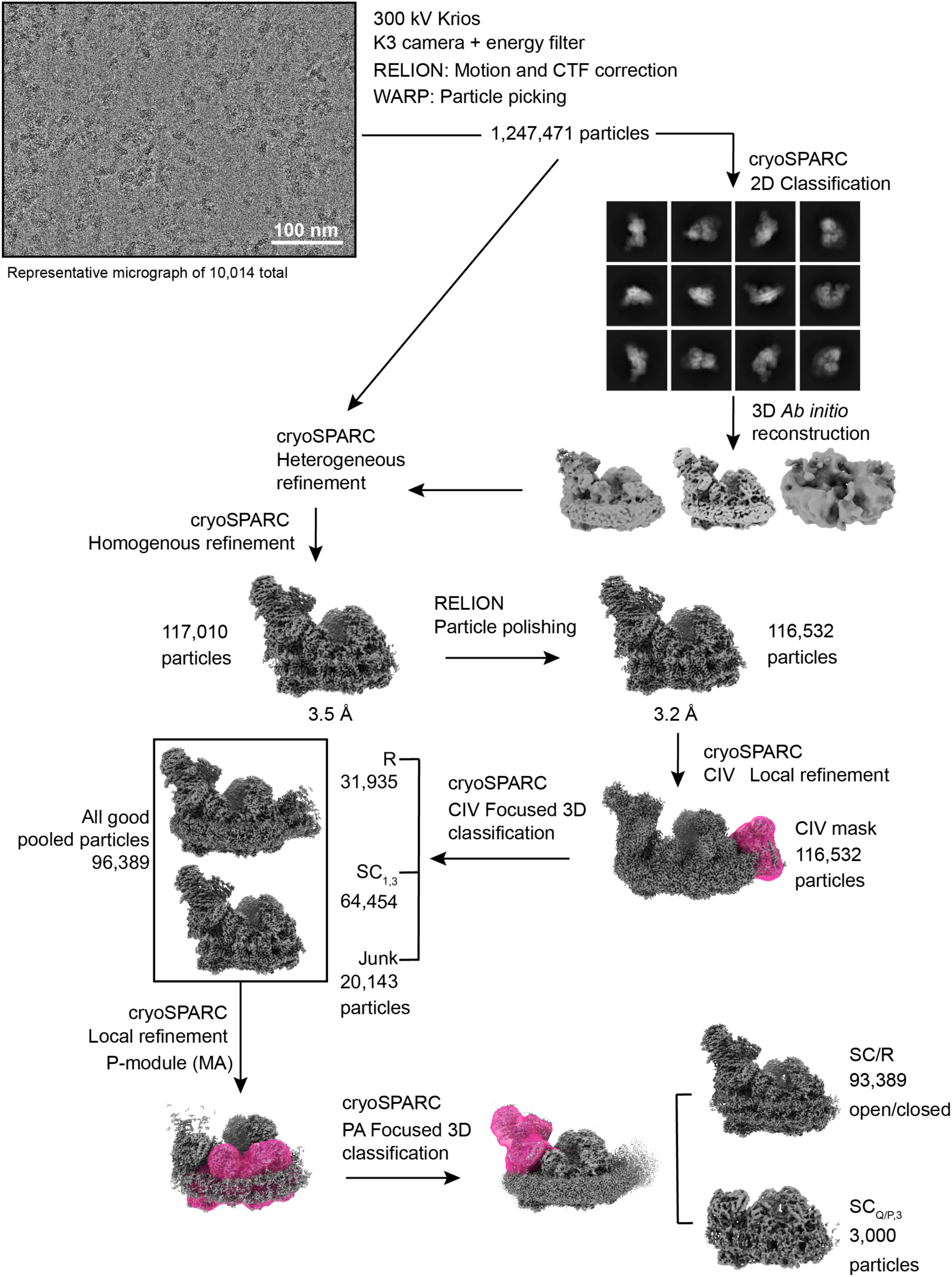
WT data processing overview. Data were collected at the SLAC-Stanford CryoEM Center on a 300 kV Titan Krios microscope (TEM-beta) with Gatam K3 camera (see also Table S1). After removal of poor-quality micrographs a total of 10,014 images were processed (representative micrograph shown), from which 1,247,471 particles were initially picked using WARP (*65*). The particles were sorted using a cryoSPARC heterogeneous refinement with 3D *ab initio* reconstructions from 2D classification as inputs. The heterogenous refinement yielded 117,010 particles that were imported into RELION. The particle set was cleaned further in RELION and after particle polishing 116,532 particles were imported back into cryoSPARC. A CIV local refinement and CIV focused 3D classification was performed to sort R, SC_1,3_ and junk particles. 96,389 good particles were obtained, and a MA local refinement was performed followed by a PA focused 3D classification to sort N-module containing from N-less particles. 3,000 SC_Q/P,3_ particles were obtained and the 93,389 SC_1,3_ and R particles were further sorted in open and closed states. R: Respirasome; SC_1,3_: Supercomplex I+III_2_; MA: membrane arm; PA: peripheral arm; SC_Q/P,3_: Supercomplex with CI Q/P intermediate plus CIII_2_ (see Table S3).

**Figure S4.**
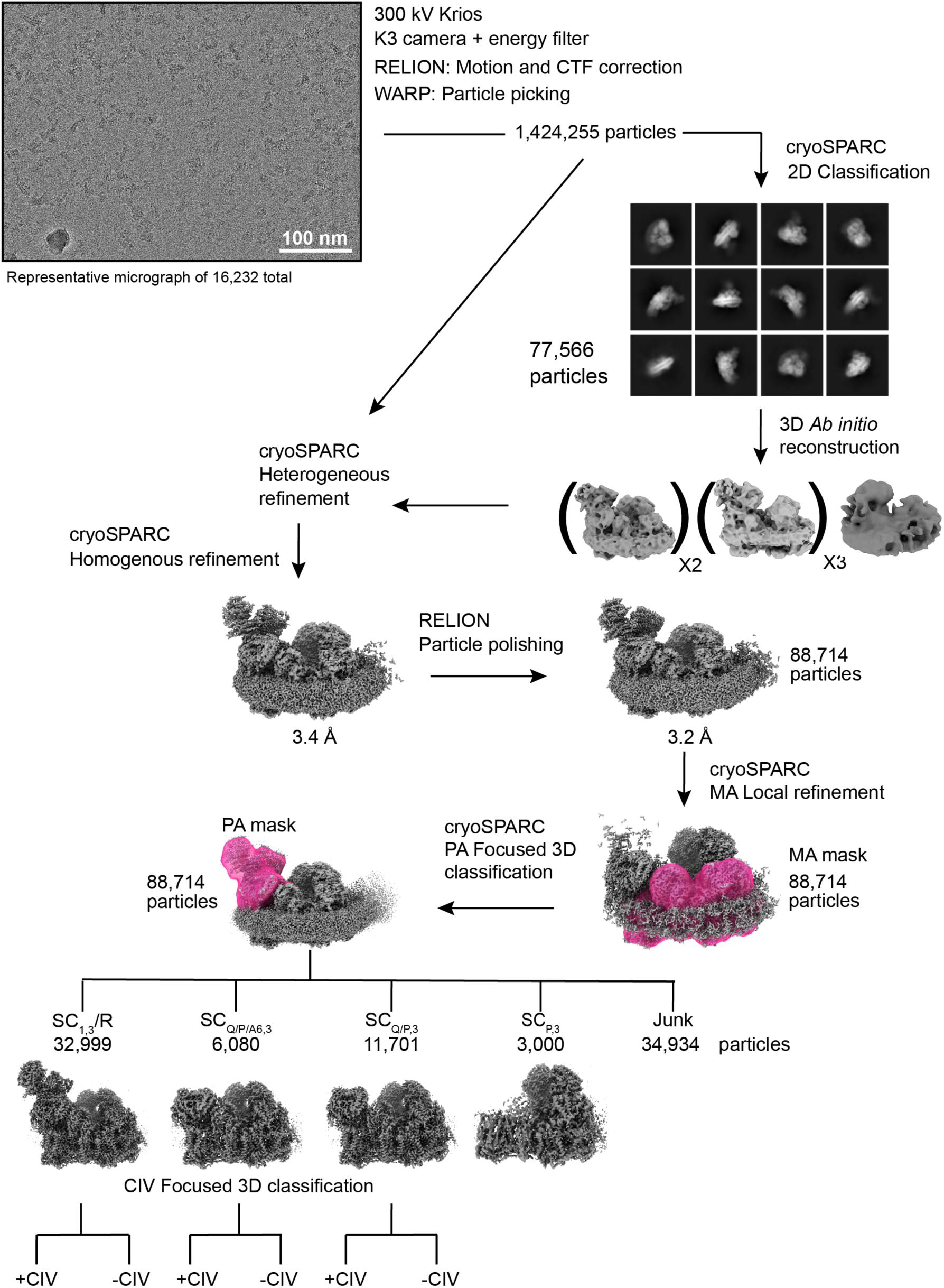
S4^KO^ data processing overview. Data were collected at the SLAC-Stanford CryoEM Center on a 300 kV Titan Krios microscope (TEM-beta) with Gatam K3 camera (see also Table S1). A total of 16,232 images were processed (representative micrograph shown), from which 1,424,255 particles were initially picked using WARP (*65*). The particles were sorted using a cryoSPARC heterogeneous refinement with 3D ab initio reconstructions from 2D classification as inputs. The heterogenous refinement yielded 95,628 particles that were imported into RELION. The particle set was cleaned further in RELION and after particle polishing 88,714 particles were imported back into cryoSPARC. A MA local refinement and PA focused 3D classification was performed to separate N-module containing, N-less particles and SC_P,3_ particles. The SC_1,3_/R, SC_Q/P/A6,3_ and SC_Q/P,3_ particle classes were further sorted via CIV focused 3D classification to find particles containing and missing CIV. R: Respirasome; SC_1,3_: Supercomplex I+III_2_; MA: membrane arm; PA: peripheral arm; SC_Q/P,3_: Supercomplex with CI Q/P intermediate plus CIII_2_, SC_Q/P/A6,3_: Supercomplex with CI Q/P intermediate with NDUFA6, NDUFAB1-α and assembly factor NDUFAF2 plus CIII_2_ (see Table S3).

**Figure S5.**
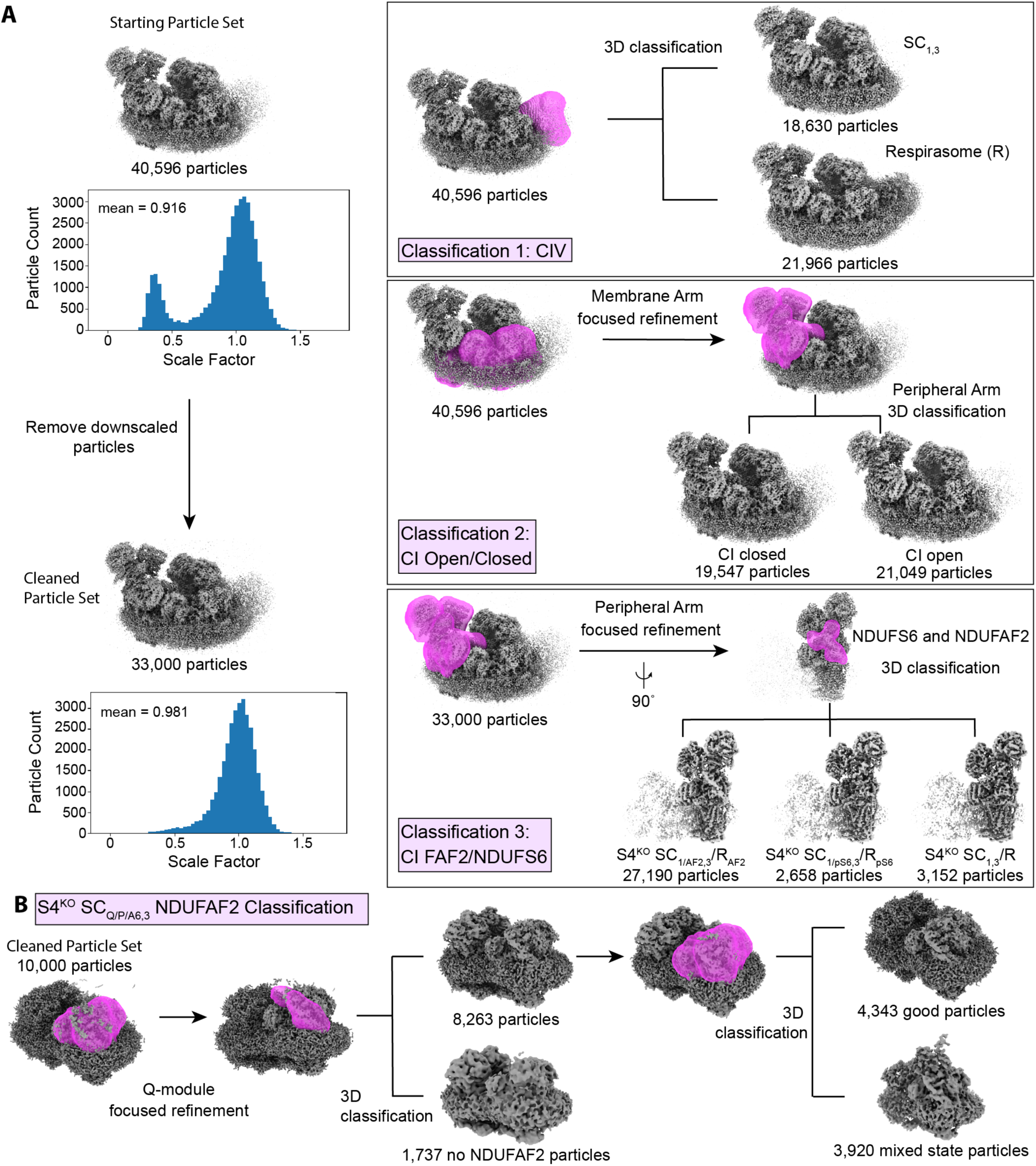
S4^KO^ additional classification strategy. **(A)** A S4^KO^ SC_1,3_/R subset of 40,596 were sorted into SC_1,3_ and R classes using a CIV focused 3D classification in cryoSPARC (classification 1). The 40,596 particles were also sorted into CI open and closed states by first doing a MA focused refinement followed by a PA 3D classification (classification 2). In parallel the 40,596 particles were cleaned by removing downscaled particles using a 3D Flex Data Prep job on cryoSPARC which yielded 33,000 good particles (left hand side). The clean set of 33,000 particles were used to sort particles into SC_1/AF2,3_, SC_1/pS6,3_ and SC_1,3_ by first doing a PA focused refinement followed by a 3D classification using a NDUFS6 and NDUFAF2 mask (Classification 3). These three classifications defined sets of particles that were compared to generate the particle classes shown in Fig. S6. For example, the union between the SC/R_1/AF2,3_, CI open and SC_1,3_ classes defines the set of SC_1/AF2,3_ open particles. **(B)** A clean set of 10,000 SC_Q/P/A6,3_ particles aligned using a Q module mask followed by a NDUFAF2 focused 3D classification. This yielded 8,263 SC_Q/P/A6,3_ particles and 1,737 SC_Q/P/A6,3 noAF2_ particles. An updated Q module mask was created and used for focused 3D classification which yielded the final SC_Q/P/A6,3_ class of 4,343 good SC_Q/P/A6,3_ particles and 3,920 junk particles. SC_1,3_: Supercomplex I+III_2_; R: Respirasome; CIV: Complex IV; CI: Complex I; MA: Membrane arm; PA: Peripheral arm; SC_1/AF2,3_: Supercomplex I+III_2_ with assembly factor NDUFAF2; SC_1/PS6,3_: Supercomplex I+III_2_ with partial NDUFS6 density; SC_Q/P,3_: Supercomplex with CI Q/P intermediate plus CIII_2_; SC_Q/P/A6,3_: Supercomplex with CI Q/P intermediate with NDUFA6, NDUFAB1-α and plus CIII_2_: SC_Q/P/A6,3 noAF2_ Supercomplex with CI Q/P intermediate with NDUFA6, NDUFAB1-α lacking NDUFAF2 and plus CIII_2_ (see Table S3).

**Figure S6.**
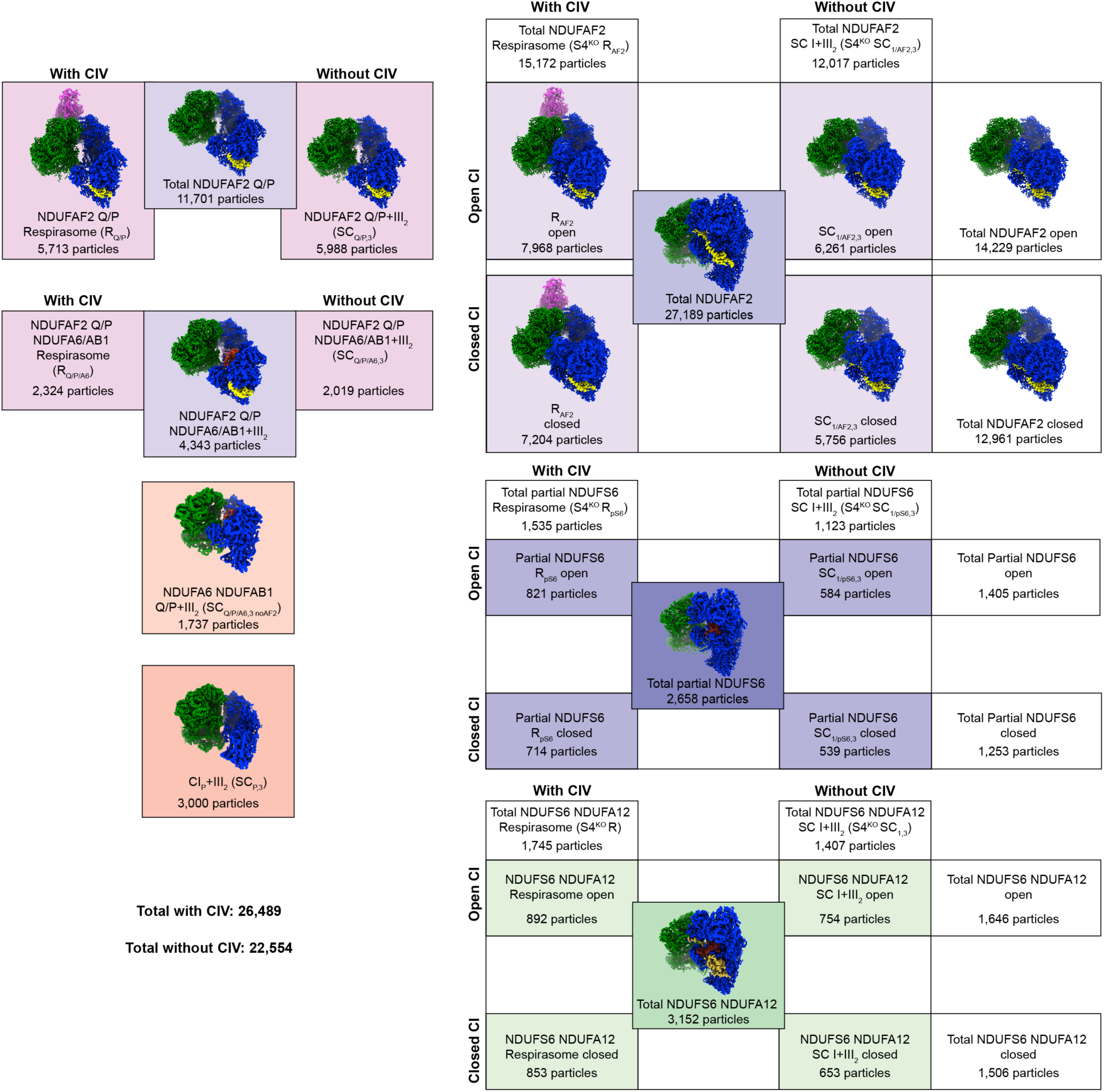
S4^KO^ structural classes and particle distribution. Diagram outlining the different structures obtained from the S4^KO^ sample through the classification strategy outlined in Fig. S5. Structures missing the N-module (R_Q/P_, SC_Q/P,3_, SC_Q/P/A6,3_, SC_Q/P/A6,3 noAF2_, SC_P,3_) are shown on the left. The SC_Q/P_ and SC_Q/P/A6,3_ classes are sorted into classes with and without CIV. Structures with the N-module (R_AF2_, SC_1/AF2,3_, R_pS6_, SC_1/pS6,3_, R and SC_1,3_) are shown on the right. These particles were sorted in open and closed states and with and without complex IV by comparison of particle sets across the multiple classifications shown in Fig. S5A. The particle number is listed for each class. Reconstructions were obtained if shown in the box. We did not obtain reconstructions for classes with less than 1,000 particles. Complexes in reconstructions are colored with CI blue, CIII_2_ green, CIV magenta, NDUFAF2 yellow, NDUFA6 red, NDUFS6 dark red and NDUFA12 mustard. SC_1,3_: Supercomplex I+III_2_; R: Respirasome; CIV: Complex IV; CI: Complex I; MA: Membrane arm; PA: Peripheral arm; SC_1/AF2,3_: Supercomplex I+III_2_ with assembly factor NDUFAF2; SC_1/pS6,3_: Supercomplex I+III_2_ with partial NDUFS6 density; SC_Q/P,3_: Supercomplex with CI Q/P intermediate plus CIII_2_; SC_Q/P/A6,3_: Supercomplex with CI Q/P intermediate with NDUFA6, NDUFAB1-α and plus CIII_2_: SC_Q/P/A6,3 noAF2_ Supercomplex with CI Q/P intermediate with NDUFA6, NDUFAB1-α lacking NDUFAF2 and plus CIII_2_ (see Table S3). S4^KO^: NDUFS*4^−/−^* (see Table S3).

**Figure S7.**
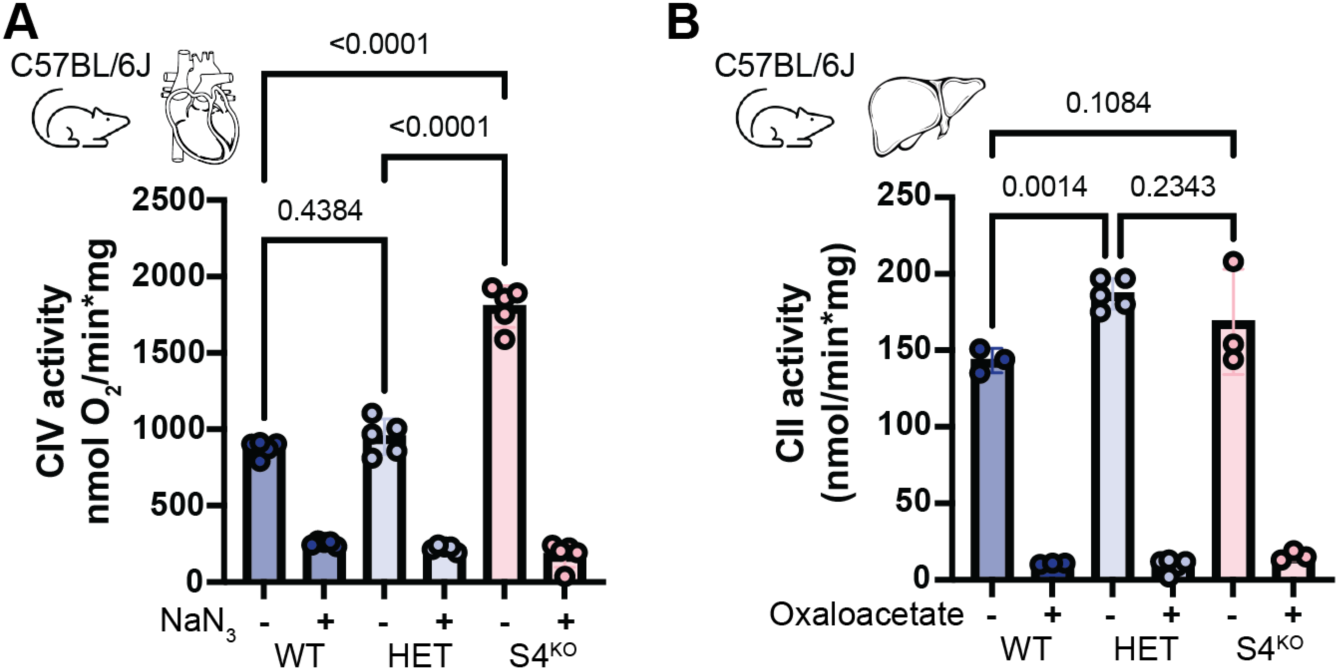
Heart CIV and liver CII functional data. (**A**) Murine heart maximal CIV oxygen consumption driven by excess ascorbate, TMPD and cyt *c*, n = 4-5, p-values from ordinary one-way ANOVA with multiple comparisons. (**B**) CII spectroscopic activity assay of murine liver mitochondrial membranes, n = 3-5, p-values from ordinary one-way ANOVA with multiple comparisons.

**Figure S8.**
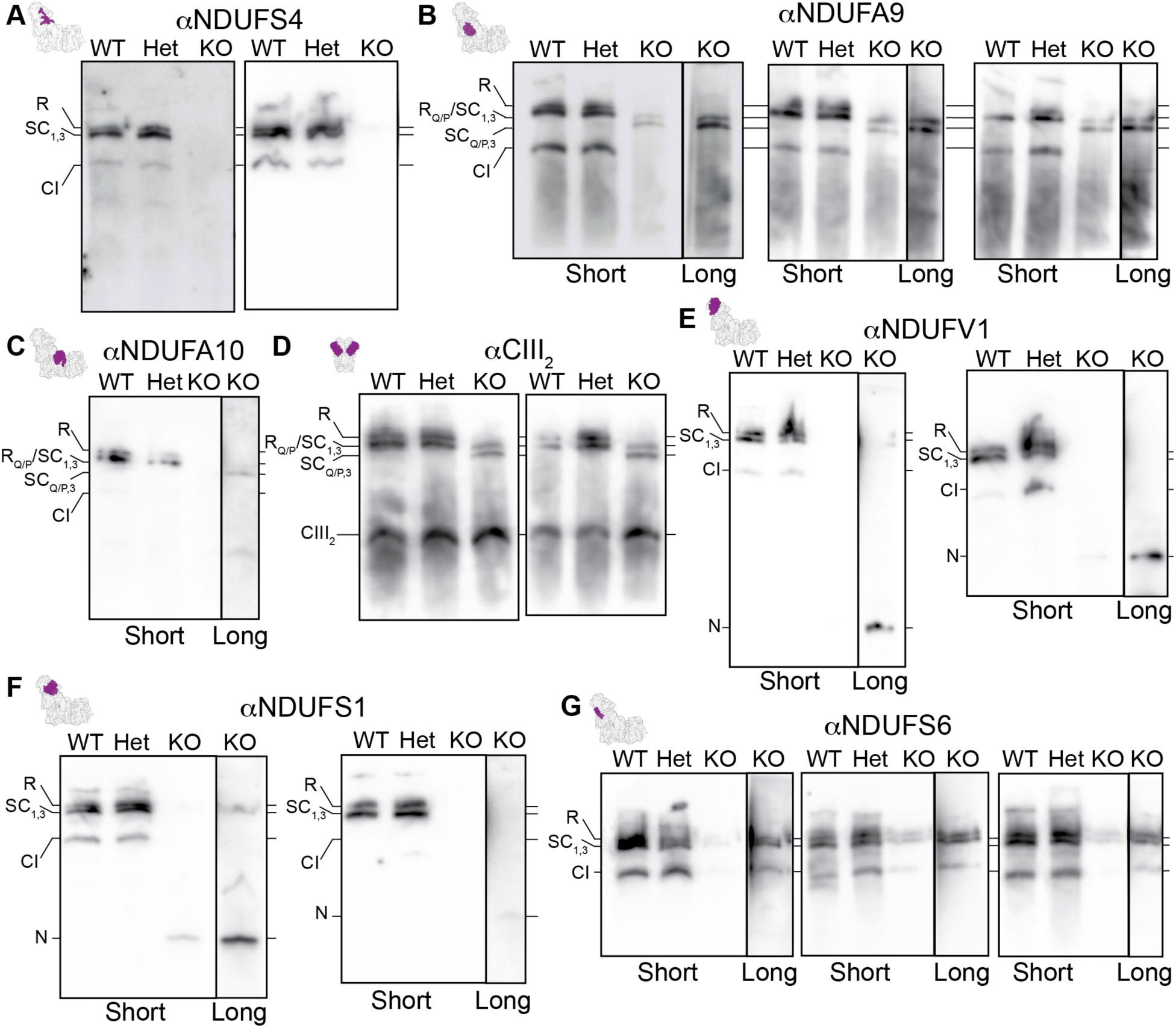
Additional Western Blots showing SCs containing CI assembly intermediates in the S4^KO^ mouse liver mitochondria. Blue Native PAGE western blots of digitonin extracted mouse liver mitochondrial complexes using primary antibodies against (**A**) CI subunit NDUFS4, (**B**) CI subunit NDUFA9, (**C**) CI subunit NDUFA10, (**D**) CIII subunit UQCRC1, (**E**) CI subunit NDUFV1, (**F**) CI subunit NDUFS1 and (**G**) CI subunit NDUFS6. The location of each subunit is indicated in purple on the structure of the complex, top left of each panel. Each blot shown is an independent repeat. Short and long labels refer to the relative exposure times. Labels: WT: NDUFS4*^+/+^*; Het: NDUFS4*^+/−^*; S4^KO^: NDUFS4*^−/−^*; R: Respirasome; SC_1,3_: Supercomplex I+III_2_; CI: complex I; R_Q/P_: Respirasome containing CI Q/P intermediate; SC_Q/P,3_: Supercomplex Complex I Q/P intermediate with CIII_2._ CIII_2_: complex III dimer; N: N-module alone.

**Figure S9.**
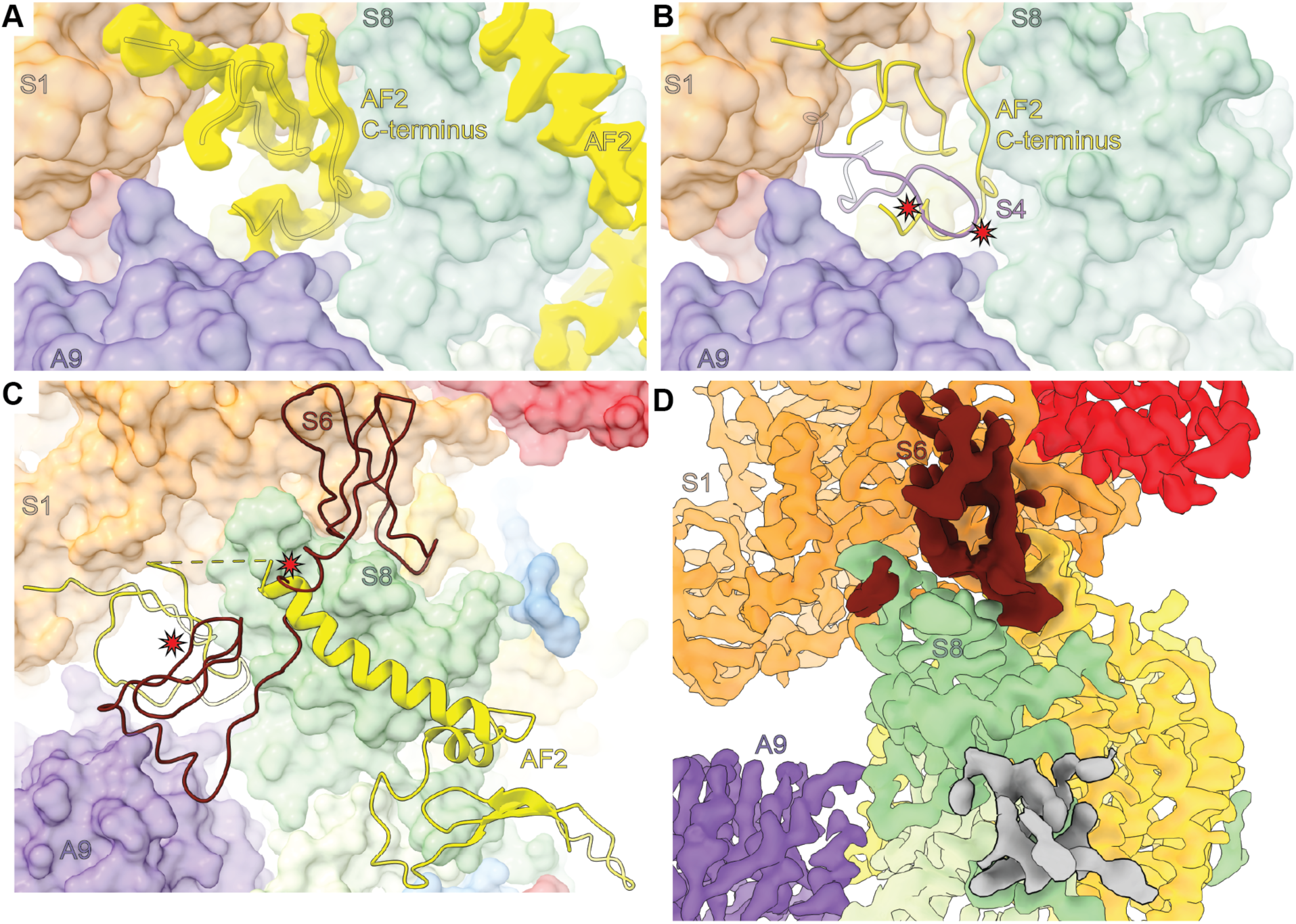
Comparison of NDUFAF2 binding to NDUFS4 and NDUFS6. **(A)** NDUFAF2 (AF2) density shown in yellow from S4^KO^ closed SC_1/AF2,3_. Density for AF2 allowed fitting of C-terminal residues AF2^P123-Y136^ residues shown in cartoon in addition to the previously modeled residues, AF2^E141-E161^ (*42*). Surfaces of NDUFS1 (S1), NDUFA9 (A9) and NDUFS8 (S8) are shown and labeled. **(B)** S4^KO^ closed SC_1/AF2,3_ structure shown with AF2 C-terminus in cartoon. NDUFS4 (S4) loop from WT closed SC_1,3_ shown as purple cartoon. Red stars indicate where the AF2 C-terminus and S4 clash. S1, A9 and S8 are shown in surface and labeled. **(C)** S4^KO^ closed SC_1/AF2,3_ structure shown with AF2 as cartoons. NDUFS6 (S6) cartoon from WT closed SC_1,3_ shown in dark red. Red stars indicate where AF2 and S6 clash. Surfaces of S1, A9 and S8 are shown and labeled. **(D)** S4^KO^ SC_1/pS6,3_ density shown colored by subunit. Partial S6 density is shown in dark red and extra density in the NDUFA12 region is shown in gray. This grey density was ambiguous and could not be modeled as either NDUFAF2 or NDUFA12 suggesting a disordered/mixed state. SC_1/AF2,3_: Supercomplex I+III_2_ with assembly factor NDUFAF2; SC_1,3_: Supercomplex I+III_2_; SC_1/PS6,3_: Supercomplex I+III_2_ with partial NDUFS6 density. WT: NDUFS4*^+/+^*; S4^KO^ NDUFS4*^−/−^* (see Table S3).

**Figure S10.**
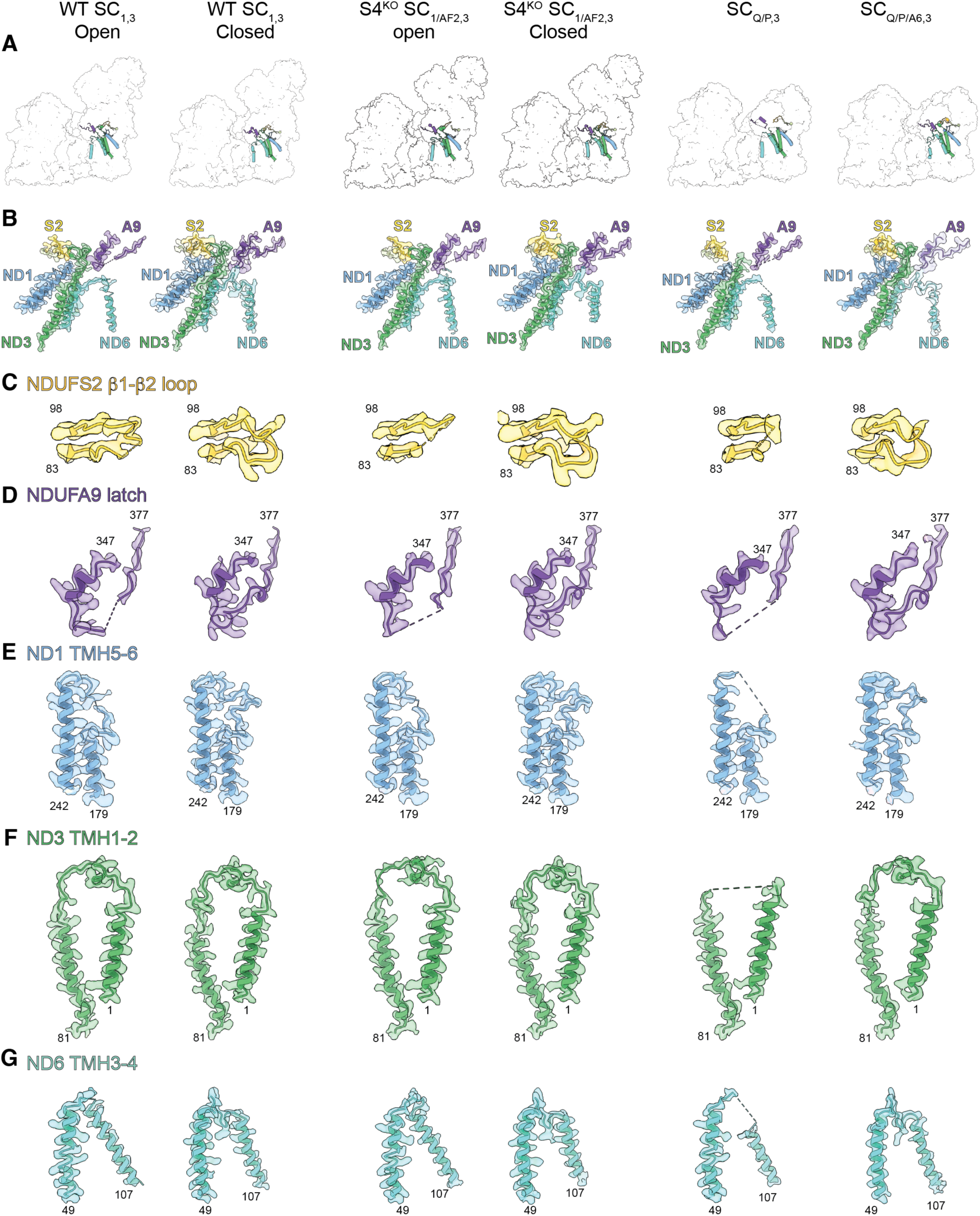
Structural features of active site loops across different states. (**A**) CI assembly for, from left to right, WT SC_1,3_ Open, WT SC_1,3_ Closed, S4^KO^ SC_1/AF2,3_ Open, S4^KO^ SC_1/AF2,3_ Closed, SC_Q/P,3_ and SC_Q/P/A6,3_ is shown as a transparent surface and the Q-site loops and the interface forming loops are show as cartoons. (**B**) Zoomed-in view of the Q-site loops and the interface forming loops for, from left to right WT SC_1,3_ Open, WT SC_1,3_ Closed, S4^KO^ SC_1/AF2,3_ Open, S4^KO^ SC_1/AF2,3_ closed, SC_Q/P,3_ and SC_Q/P/A6,3_ show as cartoons embedded in the respective cryoEM density (**C**) CryoEM density map and model of NDUFS2 β1-β2 loop (aa 83 - 98) for, from left to right, WT SC_1,3_ Open, WT SC_1,3_ Closed, S4^KO^ SC_1/AF2,3_ Open, S4^KO^ SC_1/AF2,3_ Closed, SC_Q/P,3_ and SC_Q/P/A6,3_. (**D**) CryoEM density map and model of NDUFA9 latch (aa 347 - 377) for, from left to right, WT SC_1,3_ Open, WT SC_1,3_ Closed, S4^KO^ SC_1/AF2,3_ Open, S4^KO^ SC_1/AF2,3_ Closed, SC_Q/P,3_ and SC_Q/P/A6,3_. (**E**) CryoEM density map and model of ND1 TMH5-6 (aa 179 - 242) for, from left to right, WT SC_1,3_ Open, WT SC_1,3_ Closed, S4^KO^ SC_1/AF2,3_ Open, S4^KO^ SC_1/AF2,3_ closed, SC_Q/P,3_ and SC_Q/P/A6,3_. (**F**) CryoEM density map and model of ND3 TMH1-2 (aa 1-81) for, from left to right, WT SC_1,3_ Open, WT SC_1,3_ Closed SC_1/AF2,3_, S4^KO^ SC_1/AF2,3_ Open, S4^KO^ SC_1/AF2,3_ closed, SC_Q/P,3_ and SC_Q/P/A6,3_. (**G**) CryoEM density map and model of ND6 TMH3-4 (aa 49 −107) for, from left to right, WT SC_1,3_ Open, WT SC_1,3_ Closed, S4^KO^ SC_1/AF2,3_ Open, S4^KO^ SC_1/AF2,3_ closed, SC_Q/P,3_ and SC_Q/P/A6,3_G

**Figure S11.**
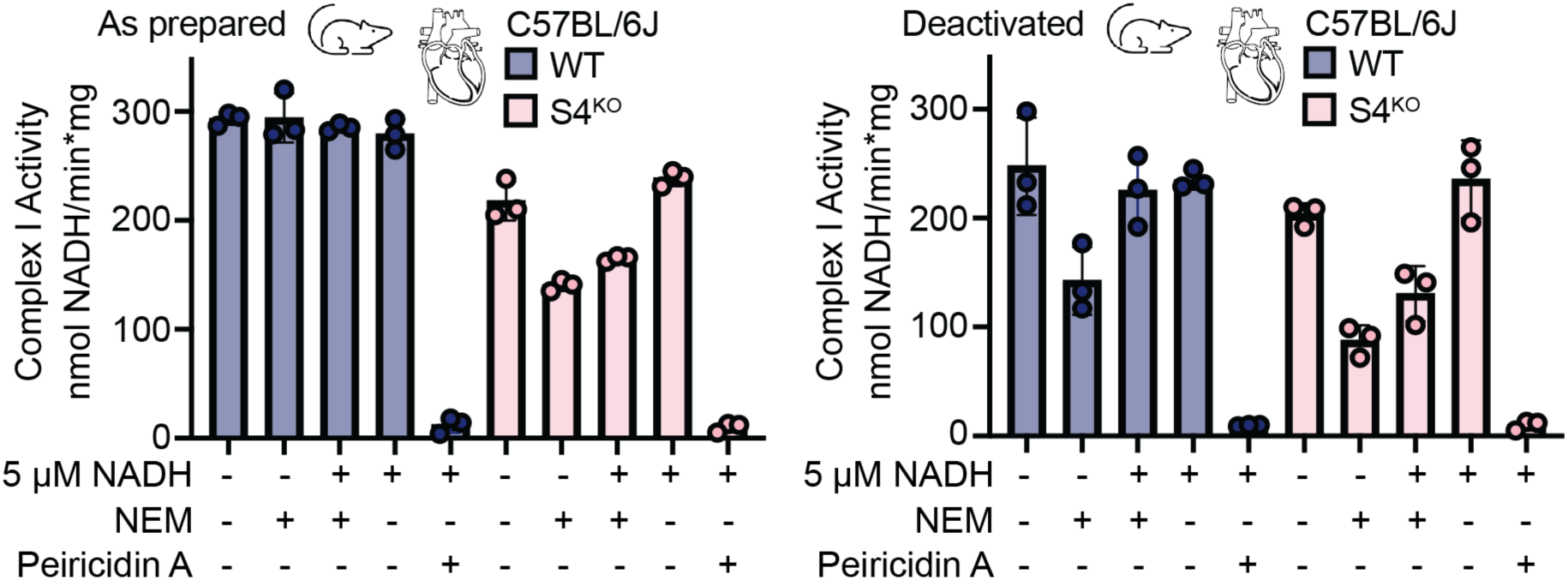
Functional characterization of the active and deactive states in the S4^KO^. Functional characterization of A-to-D transition from WT and S4^KO^ murine heart mitochondrial membranes as prepared (left) or deactivated (right) by measuring NADH oxidation at 340 nm. 2 mM N-ethylmaleimide (NEM), 5 μM NADH (pre activation) and 4 µM Piericidin A were used where indicated, n=3. WT: NDUFS4^+/+^; S4^KO^: NDUFS4^−/−^.

**Figure S12.**
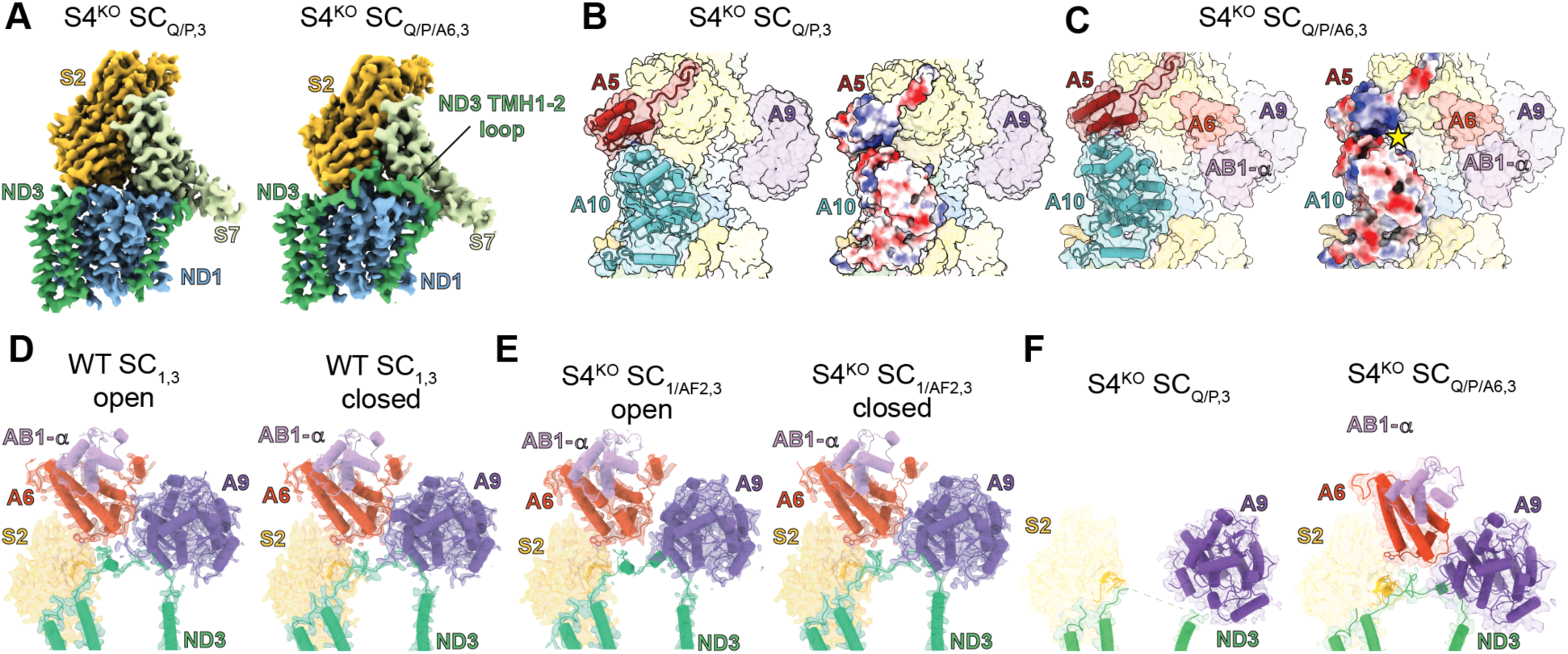
Comparison of CI_Q/P_ and CI_Q/P/A6_ structural features. **(A)** ND3 Density from S4^KO^ SC_Q/P,3_ (left) and SC_Q/P/A6,3_ (right) colored by subunit, showing the ordering of the ND3 TMH1-2 loop after NDUFA6 binding. Other nearby subunits have been removed for clarity. **(B)** S4^KO^ SC_Q/P,3_ shown in transparent surface with NDUFA5 (A5) and NDUFA10 (A10) also shown in cartoon (left). Surface electrostatics for A5 and A10 shown (right). **(C)** S4^KO^ SC_Q/P/A6,3_ shown in transparent surface with A5 and A10 also shown in cartoon (left). Surface electrostatics for A5 and A10 shown (right). The star indicates the equivalent electropositive surface that would be impacted by the NDUFA5^R126Q^ mutation in *C. elegans*. **(D)** Density from WT open SC_1,3_ (left) and closed SC_1,3_ (right) with cartoon overlay colored by subunit. **(E)** Density from S4^KO^ open SC_1/AF2,3_ (left) and S4^KO^ closed SC_1/AF2,3_ (right) with cartoon overlay colored by subunit. **(F)** Density from S4^KO^ SC_Q/P,3_ (left) and SC_Q/P/A6,3_ (right) with cartoon overlay colored by subunit. SC_Q/P,3_: Supercomplex CI Q/P intermediate plus CIII_2_; SC_Q/P/A6,3_: Supercomplex CI Q/P intermediate with NDUFA6, NDUFAB1-α plus CIII_2_; SC_1,3_: Supercomplex I+III_2_; SC_1/AF2,3_: Supercomplex I+III_2_ with assembly factor NDUFAF2. NDUFS2: mustard; ND3: green; ND1: blue; NDUFS7: light green; NDUFA6: red-orange; NDUFA9: purple; NDUFAB1-α: lavender; NDUFA5: red; NDUFA:10 turquoise. WT: NDUFS4*^+/+^*; S4^KO^: NDUFS4*^−/−^*.

**Table S1.**
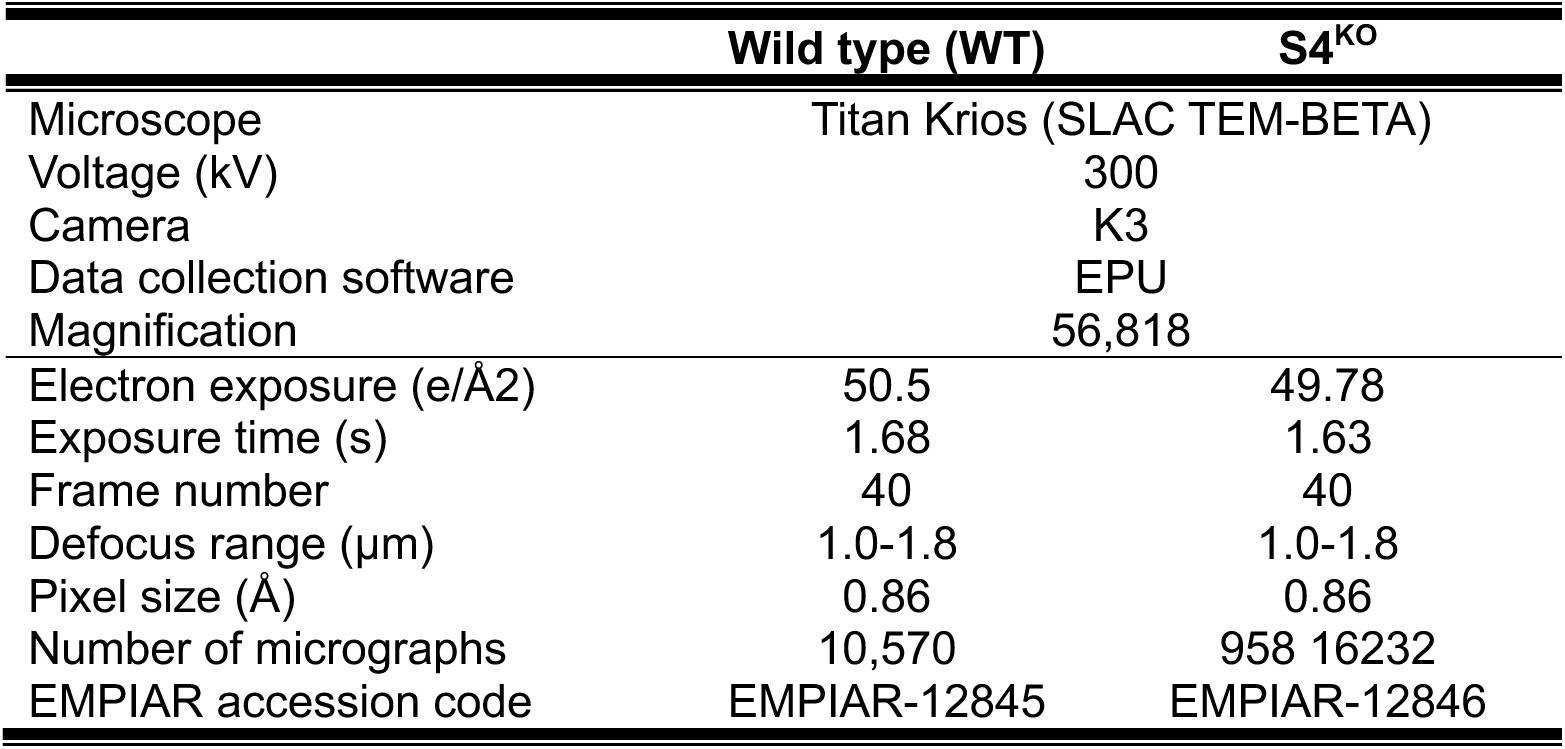
Data collection and image processing.

**Table S2.**
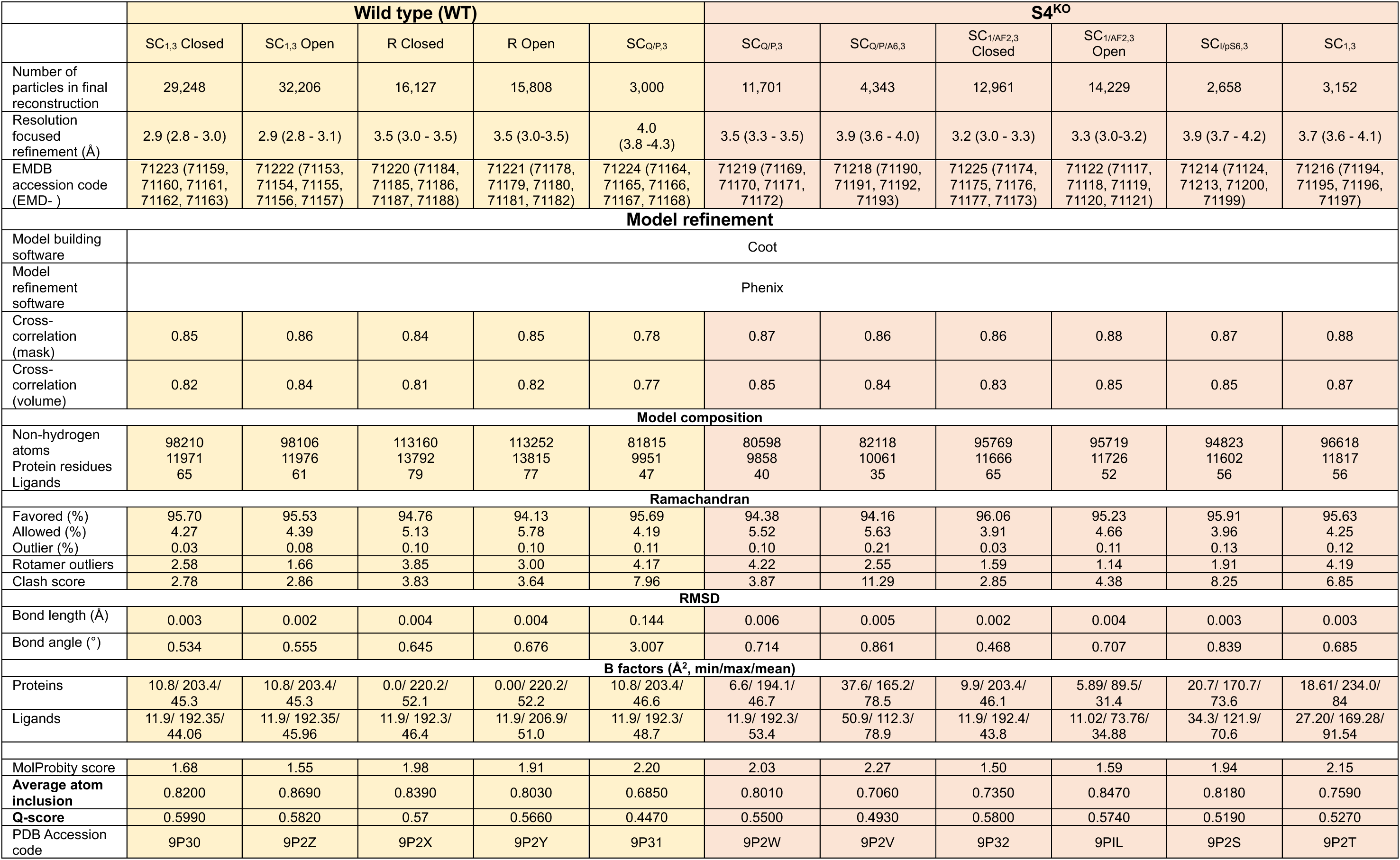
Model refinement statistics.

**Table S3.**
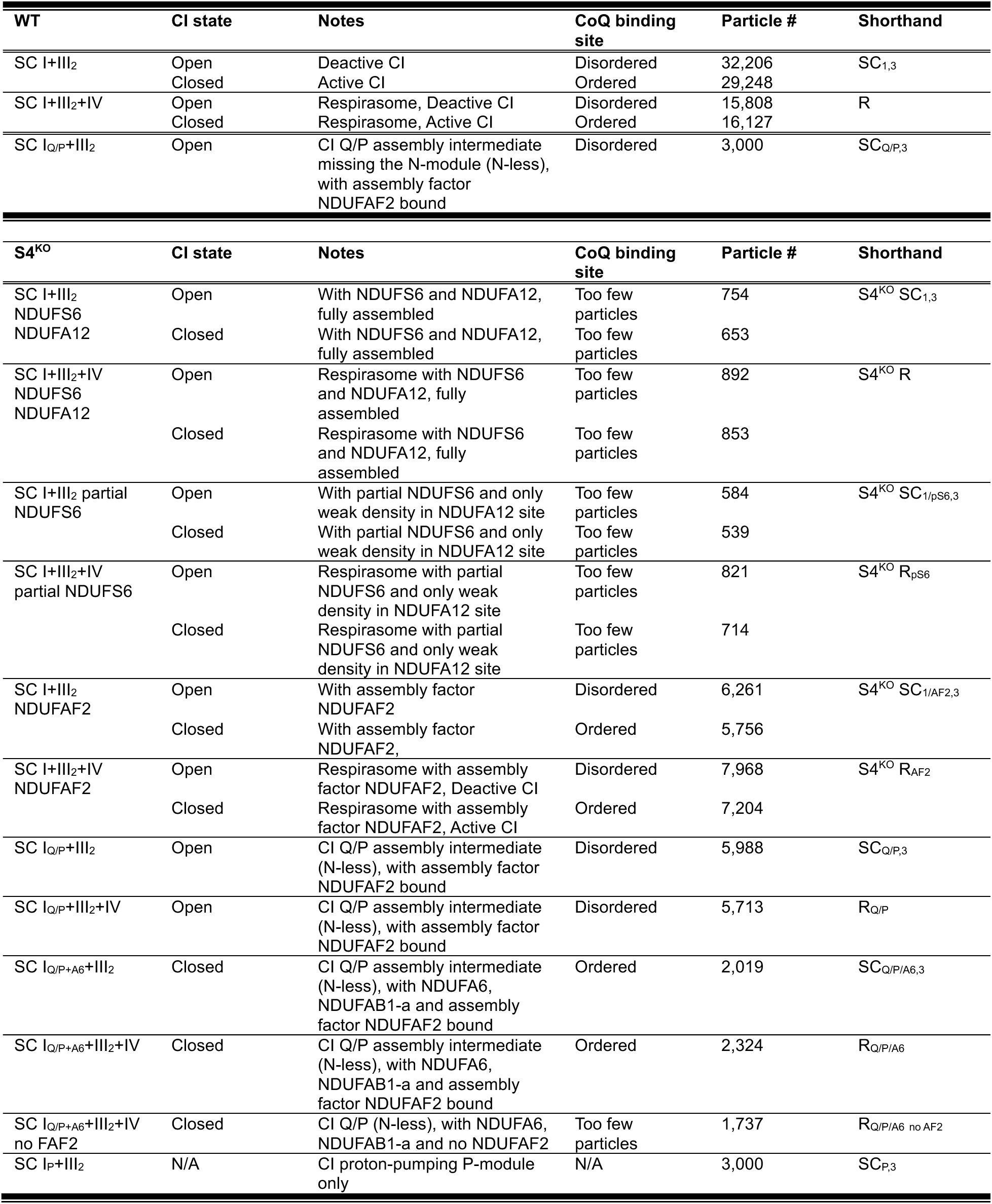
Structural states for WT and S4^KO^ liver SCs.

## References

1. V. G. Antico Arciuch, M. E. Elguero, J. J. Poderoso, M. C. Carreras, Mitochondrial regulation of cell cycle and proliferation. Antioxid Redox Signal 16, 1150–1180 (2012).

2. A. V. Kudryavtseva et al., Mitochondrial dysfunction and oxidative stress in aging and cancer. Oncotarget 7, 44879–44905 (2016).

3. L. Ernster, G. Schatz, Mitochondria: a historical review. J Cell Biol 91, 227s–255s (1981).

4. B. Morio, B. Panthu, A. Bassot, J. Rieusset, Role of mitochondria in liver metabolic health and diseases. Cell Calcium 94, 102336 (2021).

5. K. M. Davies, T. B. Blum, W. Kuhlbrandt, Conserved in situ arrangement of complex I and III(2) in mitochondrial respiratory chain supercomplexes of mammals, yeast, and plants. Proc Natl Acad Sci U S A 115, 3024–3029 (2018).

6. H. Schagger, K. Pfeiffer, Supercomplexes in the respiratory chains of yeast and mammalian mitochondria. EMBO J 19, 1777–1783 (2000).

7. W. Zheng, P. Chai, J. Zhu, K. Zhang, High-resolution in situ structures of mammalian respiratory supercomplexes. Nature 631, 232–239 (2024).

8. J. E. Walker, The NADH: ubiquinone oxidoreductase (complex I) of respiratory chains. Q Rev Biophys 25, 253–324 (1992).

9. S. Guerrero-Castillo et al., The Assembly Pathway of Mitochondrial Respiratory Chain Complex I. Cell Metab 25, 128–139 (2017).

10. I. Vercellino, L. A. Sazanov, The assembly, regulation and function of the mitochondrial respiratory chain. Nat Rev Mol Cell Biol 23, 141–161 (2022).

11. M. Protasoni et al., Respiratory supercomplexes act as a platform for complex III-mediated maturation of human mitochondrial complexes I and IV. Embo Journal 39, (2020).

12. T. Lobo-Jarne et al., Multiple pathways coordinate assembly of human mitochondrial complex IV and stabilization of respiratory supercomplexes. Embo Journal 39, (2020).

13. E. Fernandez-Vizarra, C. Ugalde, Cooperative assembly of the mitochondrial respiratory chain. Trends Biochem Sci 47, 999–1008 (2022).

14. M. A. Calvaruso et al., Mitochondrial complex III stabilizes complex I in the absence of NDUFS4 to provide partial activity. Hum Mol Genet 21, 115–120 (2012).

15. M. McKenzie, M. Lazarou, D. R. Thorburn, M. T. Ryan, Mitochondrial respiratory chain supercomplexes are destabilized in Barth Syndrome patients. J Mol Biol 361, 462–469 (2006).

16. F. Distelmaier et al., Mitochondrial complex I deficiency: from organelle dysfunction to clinical disease. Brain 132, 833–842 (2009).

17. D. M. Kirby et al., Respiratory chain complex I deficiency: an underdiagnosed energy generation disorder. Neurology 52, 1255–1264 (1999).

18. Y. Y. Ma et al., Clinical, biochemical, and genetic analysis of the mitochondrial respiratory chain complex I deficiency. Medicine (Baltimore) 97, e11606 (2018).

19. K. Fiedorczuk et al., Atomic structure of the entire mammalian mitochondrial complex I. Nature 538, 406–410 (2016).

20. M. E. Breuer, P. H. Willems, J. A. Smeitink, W. J. Koopman, M. Nooteboom, Cellular and animal models for mitochondrial complex I deficiency: a focus on the NDUFS4 subunit. IUBMB Life 65, 202–208 (2013).

21. O. M. Russell, G. S. Gorman, R. N. Lightowlers, D. M. Turnbull, Mitochondrial Diseases: Hope for the Future. Cell 181, 168–188 (2020).

22. A. E. Frazier, D. R. Thorburn, A. G. Compton, Mitochondrial energy generation disorders: genes, mechanisms, and clues to pathology. J Biol Chem 294, 5386–5395 (2019).

23. M. A. E. van de Wal et al., Ndufs4 knockout mouse models of Leigh syndrome: pathophysiology and intervention. Brain 145, 45–63 (2022).

24. S. E. Kruse et al., Mice with mitochondrial complex I deficiency develop a fatal encephalomyopathy. Cell Metab 7, 312–320 (2008).

25. M. J. W. Adjobo-Hermans et al., NDUFS4 deletion triggers loss of NDUFA12 in Ndufs4(-/-) mice and Leigh syndrome patients: A stabilizing role for NDUFAF2. Biochim Biophys Acta Bioenerg 1861, 148213 (2020).

26. F. Kahlhofer, M. Gansen, V. Zickermann, Accessory Subunits of the Matrix Arm of Mitochondrial Complex I with a Focus on Subunit NDUFS4 and Its Role in Complex I Function and Assembly. Life (Basel) 11, (2021).

27. E. Fernandez-Vizarra, V. Tiranti, M. Zeviani, Assembly of the oxidative phosphorylation system in humans: what we have learned by studying its defects. Biochim Biophys Acta 1793, 200–211 (2009).

28. C. Liang et al., Formation of I2+III2 supercomplex rescues respiratory chain defects. Cell Metabolism 37, (2025).

29. I. H. Jain et al., Hypoxia as a therapy for mitochondrial disease. Science 352, 54–61 (2016).

30. M. Ferrari et al., Hypoxia treatment reverses neurodegenerative disease in a mouse model of Leigh syndrome. Proc Natl Acad Sci U S A 114, E4241–E4250 (2017).

31. J. D. Meisel et al., Hypoxia and intra-complex genetic suppressors rescue complex I mutants by a shared mechanism. Cell 187, 659–675 e618 (2024).

32. A. A. Agip et al., Cryo-EM structures of complex I from mouse heart mitochondria in two biochemically defined states. Nat Struct Mol Biol 25, 548–556 (2018).

33. I. Vercellino, L. A. Sazanov, SCAF1 drives the compositional diversity of mammalian respirasomes. Nat Struct Mol Biol 31, 1061–1071 (2024).

34. D. N. Grba, J. J. Wright, Z. Yin, W. Fisher, J. Hirst, Molecular mechanism of the ischemia-induced regulatory switch in mammalian complex I. Science 384, 1247–1253 (2024).

35. J. S. Sousa, D. J. Mills, J. Vonck, W. Kuhlbrandt, Functional asymmetry and electron flow in the bovine respirasome. Elife 5, (2016).

36. J. A. Leks, K. Fiedorczuk, G. Degliesposti, M. Skehel, L. A. Sazanov, Structures of Respiratory Supercomplex I+III(2) Reveal Functional and Conformational Crosstalk. Mol Cell 75, 1131–1146 e1136 (2019).

37. I. Ogilvie, N. G. Kennaway, E. A. Shoubridge, A molecular chaperone for mitochondrial complex I assembly is mutated in a progressive encephalopathy. J Clin Invest 115, 2784–2792 (2005).

38. K. Parey et al., High-resolution cryo-EM structures of respiratory complex I: Mechanism, assembly, and disease. Sci Adv 5, eaax9484 (2019).

39. A. Padavannil, M. G. Ayala-Hernandez, E. A. Castellanos-Silva, J. A. Leks, The Mysterious Multitude: Structural Perspective on the Accessory Subunits of Respiratory Complex I. Front Mol Biosci 8, 798353 (2021).

40. V. Kravchuk et al., A universal coupling mechanism of respiratory complex I. Nature 609, 808–814 (2022).

41. H. Angerer et al., The LYR protein subunit NB4M/NDUFA6 of mitochondrial complex I anchors an acyl carrier protein and is essential for catalytic activity. P Natl Acad Sci USA 111, 5207–5212 (2014).

42. Z. Yin, A. A. Agip, H. R. Bridges, J. Hirst, Structural insights into respiratory complex I deficiency and assembly from the mitochondrial disease-related ndufs4(-/-) mouse. EMBO J 43, 225–249 (2024).

43. M. M. Roessler et al., Direct assignment of EPR spectra to structurally defined iron-sulfur clusters in complex I by double electron-electron resonance. Proc Natl Acad Sci U S A 107, 1930–1935 (2010).

44. D. Kampjut, L. A. Sazanov, The coupling mechanism of mammalian respiratory complex I. Science 370, (2020).

45. J. N. Blaza, K. R. Vinothkumar, J. Hirst, Structure of the Deactive State of Mammalian Respiratory Complex I. Structure 26, 312–319 e313 (2018).

46. Z. Yin et al., Structural basis for a complex I mutation that blocks pathological ROS production. Nature Communications 12, (2021).

47. E. T. Chouchani et al., Cardioprotection by S-nitrosation of a cysteine switch on mitochondrial complex I. Nat Med 19, 753–759 (2013).

48. J. J. Wright et al., Reverse Electron Transfer by Respiratory Complex I Catalyzed in a Modular Proteoliposome System. J Am Chem Soc 144, 6791–6801 (2022).

49. L. A. Sazanov, From the ‘black box’ to ‘domino effect’ mechanism: what have we learned from the structures of respiratory complex I. Biochem J 480, 319–333 (2023).

50. C. L. Alston et al., Bi-allelic Mutations in NDUFA6 Establish Its Role in Early-Onset Isolated Mitochondrial Complex I Deficiency. Am J Hum Genet 103, 592–601 (2018).

51. E. Fernandez-Vizarra et al., Two independent respiratory chains adapt OXPHOS performance to glycolytic switch. Cell Metab 34, 1792–1808 e1796 (2022).

52. I. Vercellino, L. A. Sazanov, Structure and assembly of the mammalian mitochondrial supercomplex CIII(2)CIV. Nature 598, 364–367 (2021).

53. I. Chung, D. N. Grba, J. J. Wright, J. Hirst, Making the leap from structure to mechanism: are the open states of mammalian complex I iden1fied by cryoEM resting states or catalytic intermediates? Curr Opin Struct Biol 77, 102447 (2022).

54. D. Kampjut, L. A. Sazanov, Structure of respiratory complex I - An emerging blueprint for the mechanism. Curr Opin Struc Biol 74, (2022).

55. A. N. A. Agip, J. N. Blaza, J. G. Fedor, J. Hirst, Mammalian Respiratory Complex I Through the Lens of Cryo-EM. Annu Rev Biophys 48, 165–184 (2019).

56. L. Kussmaul, J. Hirst, The mechanism of superoxide production by NADH:ubiquinone oxidoreductase (complex I) from bovine heart mitochondria. Proc Natl Acad Sci U S A 103, 7607–7612 (2006).

57. R. L. S. Goncalves et al., CoQ imbalance drives reverse electron transport to disrupt liver metabolism. Nature, (2025).

58. E. T. Chouchani et al., Ischaemic accumulation of succinate controls reperfusion injury through mitochondrial ROS. Nature 515, 431-+ (2014).

59. H. R. Bridges, E. Bill, J. Hirst, Mossbauer Spectroscopy on Respiratory Complex I: The Iron-Sulfur Cluster Ensemble in the NADH-Reduced Enzyme Is Partially Oxidized. Biochemistry-Us 51, 149–158 (2012).

60. J. Hirst, Mitochondrial complex I. Annu Rev Biochem 82, 551–575 (2013).

61. P. Mitchell, Chemiosmotic coupling in oxidative and photosynthetic phosphorylation. Biol Rev Camb Philos Soc 41, 445–502 (1966).

62. D. G. Nicholls, S. J. Ferguson, THE CHEMIOSMOTIC PROTON CIRCUIT IN ISOLATED ORGANELLES Theory and Practice. Bioenergetics 4, 53–87 (2013).

63. E. Galemou Yoga et al., Essential role of accessory subunit LYRM6 in the mechanism of mitochondrial complex I. Nat Commun 11, 6008 (2020).

64. E. Fernandez-Vizarra et al., Isolation of mitochondria for biogenetical studies: An update. Mitochondrion 10, 253–262 (2010).

65. D. Tegunov, P. Cramer, Real-time cryo-electron microscopy data preprocessing with Warp. Nat Methods 16, 1146–1152 (2019).

